# Rei1 and Reh1 facilitate the loading of eL24

**DOI:** 10.64898/2026.03.31.715693

**Authors:** Ran Lin, Madison J. Reynolds, Nila R. Shankar, Arlen W. Johnson

**Author notes:** Full mailing addresses and zip codes for all authors: 2506 Speedway, NMS 1.246, Molecular Biosciences Department, The University of Texas at Austin, Austin, TX 78712-1095, U.S.A.

## Abstract

The correct assembly of ribosomes is essential for viability and faithful gene expression. In eukaryotic cells, the pre-40S and pre-60S ribosomal subunits are largely pre-assembled in the nucleolus before they are exported to the cytoplasm for final maturation. Although most ribosomal proteins of the large subunit are loaded onto pre-60S particles in the early nucleolar steps, a few, including eL24, are loaded in the cytoplasm. eL24 is thought to recruit the zinc-finger protein Rei1 (ZNF622 in humans). In yeast, Rei1 has a paralog, Reh1. While we and others have previously shown that Rei1 facilitates the removal of Arx1, Rei1 and Reh1 appear to have an additional unknown function. To identify this function, we first examined the protein composition of pre-60S subunits isolated from *rei1Δ reh1Δ* mutant cells and found that these subunits were specifically defective for eL24. However, the absence of eL24 did not impair Rei1 binding to pre-60S. Moreover, overexpression of eL24 suppressed the growth defect of the double mutant. As an alternative approach to understanding the function of Rei1 and Reh1, we screened for bypass suppressors of the growth defect of *rei1Δ reh1Δ* cells. We identified mutations in the genes coding for ribosomal protein uL3, the GTPase Lsg1 and the protein phosphatase Ppq1. Importantly, these suppressors all partially reversed the eL24 loading defect of *rei1Δ reh1Δ* cells. Based on these results, we propose a revised order of cytoplasmic assembly events where Rei1 and Reh1 facilitate the recruitment of eL24 to the pre-60S particle.

## Introduction

Ribosomes are responsible for the rapid and accurate translation of our genetic code into proteins, a highly dynamic process that depends on the exquisitely complex structure of the ribosome. Each of the two ribosomal subunits has a distinct role in translation: the large (60S in eukaryotes) subunit carries out polypeptide synthesis while the small (40S) subunit decodes mRNA. Eukaryotic ribosome biogenesis requires more than 200 trans-acting factors for the complex events of pre-rRNA cleavage and modification, RNA folding and protein assembly (Woolford 2015; Baßler and Hurt 2019; Bohnsack and Bohnsack 2019; Broeck and Klinge 2024). Biogenesis factors also direct nuclear export and cytoplasmic maturation to produce functional ribosomes (Ho et al. 2000; Gadal et al. 2001; W. Yao et al. 2007; Y. Yao et al. 2010; Bradatsch et al. 2012). Although the two subunits are derived from the same precursor transcript, early processing steps separate the two pre-subunits which are then independently matured and exported (Kressler et al. 2017). At the time of their export, the two subunits are largely preassembled. However, several ribosomal proteins (rproteins) still need to be assembled and positioned correcly in the cytoplasm including uL10, uL11, uL16, eL24, eL40 and eL41 (Saveanu et al. 2003; Kappel et al. 2012; Ma et al. 2017; Konikkat and Woolford, 2017; Kargas et al. 2019). In addition, several assembly factors must be removed (Lo et al. 2010). Upon export, the type II AAA-ATPase Drg1 initiates cytoplasmic maturation of the pre-60S subunit by releasing the placeholder protein Rlp24 (Kappel et al. 2012; Loibl et al. 2014; Ma et al. 2022). Rlp24 shuttles between the nucleus and the cytoplasm and plays an essential role during the ribosome biogenesis. The N-terminal domain of eL24 is homologous to the N-terminal domain of Rlp24, and once Rlp24 is released, the binding site for eL24 is available (Saveanu et al. 2003).

The eukaryotic rprotein eL24 is one of 14 nonessential ribosomal proteins (Steffen et al. 2012). It contains 155 amino acid residues and in yeast it is encoded by two paralogous genes *RPL24A* and *RPL24B*. The encoded proteins differ by only five amino acids. eL24 is composed of an N-terminal domain, a linker region and a C-terminal alpha-helix (Ben-Shem et al. 2011). It performs a number of tasks required for optimal ribosome functionality including forming the eB13 and B6 intersubunit bridges (Ben-Shem et al. 2011) and the N-terminus plays a non-essential role in translation initiation (Kisly et al. 2019). The loss of eL24 is also reported to affect translation elongation on a polyU message *in vitro* (Dresios et al. 2000, 2001).

During ribosome biogenesis, eL24 is thought to recruit Rei1 (Lebreton et al. 2006). Rei1 (ZNF622 in humans) is a zinc-finger protein that, along with the Hsp70 chaperones Ssa1/2, promotes the release of the assembly factor Arx1 and its partner Alb1 (Hung and Johnson 2006; Lebreton et al. 2006; Demoinet et al. 2007; Meyer et al. 2010). Incorporation of uL16 completes the peptidyl transferase center, together with the GTPase Lsg1 (Zhou et al. 2019), triggers the release of Nmd3 (Kargas et al. 2019). A second GTPase, Efl1, is required for the release of Tif6, which ultimately licenses the subunit for translation (Weis et al. 2015). In yeast, Rei1 has a paralog, Reh1, which replaces Rei1 at the time of Arx1 release (Kargas et al. 2019). We have recently shown that Reh1 is the last assembly factor to be released from the subunit and it is released during early rounds of translation elongation (Musalgaonkar et al. 2025). Reh1 has an additional function distinct from Rei1 in promoting the turnover of pre-60S subunits containing mutant uL16 (Chitale et al. 2025). As the primary function of Rei1 appears to be the release of Arx1, the growth defect of a *rei1*Δ mutant can be partially suppressed by the deletion of *ARX1* (Hung and Johnson 2006; Lebreton et al. 2006) and can be nearly fully suppressed by an *arx1* mutant (*arx1-S347P*) which has reduced affinity for pre-60S (Lo et al. 2010). However, the simultaneous deletion of *REI1* and *REH1* leads to a severe growth defect which cannot be suppressed by deletion of *ARX1*, suggesting that Rei1 and Reh1 have a role in addition to the release of Arx1 (Parnell and Bass 2009). Here, we have addressed this additional function of Rei1 and Reh1 and show that Rei1 and Reh1 facilitate the loading of eL24 into the pre-60S subunit.

## Materials and methods

### Strains and Plasmids

All strains are listed in Table 1. Strain AJY1544 was made by crossing *rpl24a::kanMX* and *rpl24b::kanMX*. *ybl028cΔ* (AJY4875) was generated by amplifying pFA6a-His3MX6 with AJO4440x4441 and integration into BY4741. Strains related to *YBL028C* were generated by crossing AJY4875 with appropriate *rei1Δ* and *reh1Δ* strains. AJY4700 was generated by amplifying *LSG1-TAP:HISMX6* from AJY1872 with AJO(2662x1752) and integrated into AJY4686. AJY4809 was generated by amplifying pFA6a-3HA-His3MX6 with AJO(4323x4324) and integration into BY4741. AJY4810 was generated by amplifying *LSG1-TAP:HIS* from pFA6a-3HA-His3MX6 with AJO(4323x4324) and integration into AJY4686. AJY4841(*lsg1-H621L*), AJY4842(*rpl3-V57G*), AJY4843(*lsg1-K619Q*) and AJY4844(*ppq1-G491C*) were isolated as spontaneous growth suppressors of AJY3027. AJY4876 was generated by amplifying 3xHA:His3MX6 from *pFA6a-3HA:His3MX6* with AJO(4323x4324) and integration into AJY3027. AJY4877 was generated by amplifying 3xHA:His3MX6 from *pFA6a-3HA:His3MX6* with AJO(4323x4324) and integration into AJY4844. AJY4881 was generated by amplifying *LSG1-K621L-TAP-HIS3MX* from AJY4700 using AJO(4434x1752) and integration into AJY4686. AJY4882 was generated by amplifying *LSG1-K619Q-TAP-HIS3MX* from AJY4700 using AJO(4435x1752) and integration into AJY4686. AJY4883 was generated by amplifying *LSG1-Δ607-640-TAP-HIS3MX* from AJY4700 using AJO(4436x1752) and integrated into AJY4686. AJY4884 was generated by amplifying *3xHA:His3MX6* from *pFA6a-3HA:His3MX6* with AJO(4323x4324) and integration into AJY4844. AJY4885 was made in AJY4700 by CRISPR with pAJ5625 and AJO4439 as repair template. AJY4886 was made in AJY4686 by CRISPR with pAJ5625 and AJO4439 as a repair template. AJY4887 was made in AJY3027 by CRISPR with pAJ5625 and AJO4439 as a repair template. AJY4888 was made in AJY4700 by CRISPR with pAJ5626 and AJO4442 as a repair template. AJY4889 was made in AJY4686 by CRISPR with pAJ5625 and AJO4442 as a repair template. AJY4890 was made in AJY3027 by CRISPR with pAJ5625 and AJO4442 as a repair template. AJY4857 was made by amplifying *rim15Δ::kanMX* with AJO(4387XAJO4388) from the heterozygous diploid collection (Research Genetics), and integration into BY4742. AJY4852 was made by amplifying *kanMX* with AJO(4375XAJO4376) and integration into BY4741. *NatMX* was amplified with AJO(2957 x AJO2958) from p4339 and integrated into AJY4852 to generate AJY4854. AJY4859 was made by amplifying *kanMX::NatMX* with AJO2957XAJO2958 from p4339 integrated into AJY4857. AJY4856 was made by amplifying *NatMX* with AJO(4377XAJO4378) and integrated into AJY4686. AJY4862 and AJY4863 were generated by crossing AJY1948 and AJY4860. Strains AJY5054, AJY5055, AJY5056 and AJY5057 were made by amplifying *LSG1- TAP:HIS3MX6* from AJY4700 with oligonucleotide pairs AJO(4748XAJO4376), AJO(4749XAJO4376), AJO(4750XAJO4376) and AJO(4751XAJO4376), respectively, and integration into AJY4686. Mutations were confirmed by Sanger sequencing.

**Table 1.** Strains used in this work.

| **Strains** | **Description** | **Source** |
| --- | --- | --- |
| **AJY1544** | *MATa rpl24aΔ::KanMX rpl24bΔ::KanMX his3Δ1 leu2Δ0 ura3Δ0 lys2Δ0 met15Δ0* | This work |
| **AJY1872** | *MATa LSG1-TAP:HIS3MX6 his3Δ1 leu2Δ0 ura3Δ0 met15Δ0* | Open Biosystems. |
| **AJY1948** | *MATa ARX1-GFP::HIS3MX his3Δ1 leu2Δ0 ura3Δ0 met15Δ0* | (Lo et al. 2010) |
| **AJY3027** | *MATalpha reh1Δ::KanMX rei1ΔKanMX arx1Δ::NatMX his3Δ1 leu2Δ0 ura3Δ0* | (Musalgaonkar et al. 2023) |
| **AJY3827** | *MATalpha NatMX-PGal1-3xHA-LSG1 his3Δ1 leu2Δ0 ura3Δ0* | (Malyutin et al. 2017) |
| **AJY4686** | *MATa reh1Δ::KanMX rei1ΔKanMX arx1-S347P his3Δ1 leu2Δ0 ura3Δ0* | (Musalgaonkar et al. 2023) |
| **AJY4700** | *MATa rei1Δ::KanMX reh1Δ::KanMX arx1-S347P LSG1-TAP:HIS3MX6 his3Δ1 leu2Δ0 ura3Δ0* | This work |
| **AJY4809** | *MATa RPL24A-3XHA:HIS3MX6 his3Δ1 leu2Δ0 ura3Δ0 met15Δ0* | This work |
| **AJY4810** | *MATa RPL24A-3XHA:HIS3MX6 rei1Δ::kanMX reh1Δ::kanMX arx1-S347P I his3Δ1 leu2Δ0 ura3Δ0* | This work |
| **AJY4841** | *MATalpha lsg1-H621L rei1Δ::KanMX arx1Δ::NatMXr reh1Δ::KanMX* | This work |
| **AJY4842** | *MATalpha rpl3-V57G rei1Δ::KanMX arx1Δ::NatMXr reh1Δ::KanMX* | This work |
| **AJY4843** | *MATalpha lsg1-K619Q rei1Δ::KanMX arx1Δ::NatMXr reh1Δ::KanMX* | This work |
| **AJY4844** | *MATalpha ppq1(G491C) rei1Δ::KanMX arx1Δ::NatMXr reh1Δ::KanMX* | This work |
| **AJY4852** | *MATa ppq1Δ::kanMX his3Δ1 leu2Δ0 ura3Δ0 lys2Δ0 met15Δ0* | This work |
| **AJY4854** | *MAT a ppq1Δ::kanMX::NatMX his3Δ1 leu2Δ0 met15Δ0 ura3Δ0* | This work |
| **AJY4856** | *MATa ppq1Δ::kanMX::NatMX cloNAT rei1Δ::kanMX reh1::kanMX arx1-S347P his3Δ1 leu2Δ0 ura3Δ0 lys2Δ0 met15Δ0* | This work |
| **AJY4857** | *MAT alpha rim15Δ::kanMX his3Δ1 leu2Δ0 ura3Δ0 lys2Δ0* | This work |
| **AJY4859** | *MATalpha rim15Δ::kanMX::NatMX his3Δ1 leu2Δ0 ura3Δ0 lys2Δ0* | This work |
| **AJY4860** | *MATalpha rim15Δ::NatMX rei1Δ::kanMX reh1Δ::kanMX arx1-S347P his3Δ1 leu2Δ0 ura3Δ0 lys2Δ0* | This work |
| **AJY4862** | *MATalpha arx1-S347P* | This work |
| **AJY4863** | *MATa rim15Δ::NatMX arx1-S347P* | This work |
| **AJY4875** | *MATa ybl028cΔ::HIS3MX6 his3Δ1 leu2Δ0 ura3Δ0 met15Δ0* | This work |
| **AJY4876** | *MATalpha rei1Δ::KanMX arx1Δ::NatMXr reh1Δ::KanMX RPL24A-3HA:HIS3MX6* | This work |
| **AJY4877** | *MATalpha lsg1(H621L) rei1Δ::KanMX arx1Δ::NatMXr reh1Δ::KanMX RPL24A-3HA:HIS3MX6* | This work |
| **AJY4878** | *MATalpha rpl3(V57g) rei1Δ::KanMX arx1Δ::NatMXr reh1Δ::KanMX RPL24A-3HA:HIS3MX6* | This work |
| **AJY4881** | *MATa rei1Δ::KanMX reh1Δ::KanMX arx1-S347P lsg1-H621L-TAP-HIS3MX6* | This work |
| **AJY4882** | *MATa rei1Δ::KanMX reh1Δ::KanMX arx1-S347P lsg1-K619Q-TAP-HIS3MX6* | This work |
| **AJY4883** | *MATa rei1Δ::KanMX reh1Δ::KanMX arx1-S347P lsg1-Δ607-640-TAP-HIS3MX6* | This work |
| **AJY4884** | *MATalpha ppq1(G491C) rei1Δ::KanMX arx1Δ::NatMXr reh1Δ::KanMX Rpl24A-3HA: :HISMX6 his3Δ1 leu2Δ0 ura3Δ0* | This work |
| **AJY4885** | *MATa rpl3(V57G) rei1Δ::KanMX reh1Δ::KanMX arx1-S347P LSG1-TAP:HIS3MX6 his3Δ1 leu2Δ0 ura3Δ0* | This work |
| **AJY4886** | *MATa rpl3(V57G) rei1Δ::KanMX reh1Δ::KanMX arx1-S347P* | This work |
| **AJY4887** | *MATa rpl3(V57G) rei1Δ::KanMX reh1Δ::KanMX arx1Δ::NatMX* | This work |
| **AJY4888** | *MATa ppq1(G491C) rei1Δ::KanMX reh1Δ::KanMX arx1-S347P LSG1-TAP:HIS3MX6* | This work |
| **AJY4889** | *MATa ppq1(G491C) rei1Δ::KanMX reh1Δ::KanMX arx1-S347P* | This work |
| **AJY4890** | *MATa ppq1(G491C) rei1Δ::KanMX reh1Δ::KanMX arx1Δ::NatMX* | This work |
| **AJY5054** | *MATa lsg1-T607E-TAP-HIS3MX6 rei1Δ::KanMX reh1Δ::KanMX arx1-S347P* | This work |
| **AJY5055** | *MATa lsg1-T607A-TAP-HIS3MX6 rei1Δ::KanMX reh1Δ::KanMX arx1-S347P* | This work |
| **AJY5056** | *MATa lsg1-S632E-TAP-HIS3MX6 rei1Δ::KanMX reh1Δ::KanMX arx1-S347P* | This work |
| **AJY5057** | *MATa lsg1-S632A-TAP-HIS3MX6 rei1Δ::KanMX reh1Δ::KanMX arx1-S347P* | This work |
| **BY4741** | *MATa his3Δ1 leu2Δ0 ura3Δ met15Δ* | (Baker Brachmann et al. 1998) |
| **BY4742** | *MATalpha his3Δ1 leu2Δ0 ura3Δ met15Δ* | (Baker Brachmann et al. 1998) |

All plasmids are listed in Table 2. pAJ3804 was made by inverse PCR of pAJ2520 with AJO(2438x2439). pAJ4963 was made through Golden Gate assembly with pAJ5107 and pAJ5053, using *RPL24A* amplified from gDNA with AJO(4310x4311). pAJ4964 was made through Golden Gate assembly with pAJ5104 and pAJ5053, using RPL24A amplified from gDNA with AJO(4310x4311). pAJ5460 was made by inverse PCR from pMP001 amplified with AJO(4394x4395) and followed up with BsaI ligation. pAJ5613 was made by Golden Gate assembly of pAJ5103, PCR product of gDNA YBL028C with AJO(4472x4473) and pAJ5053 with a terminator sequence ADH1 (tADH1). pAJ5614 was made by Golden Gate assembly of pAJ5106, PCR product of gDNA YBL028C with ybl028c fwdxrev and pAJ5053 with a terminator sequence ADH1 (tADH1). pAJ5625 was generated by annealing AJO4437 and AJO4438 and assembly into pAJ4247 with BsmBI. pAJ5626 was made by AJO4440 and 4441 annealed and assembled into pAJ4247 with BsmBI.

**Table 2.**
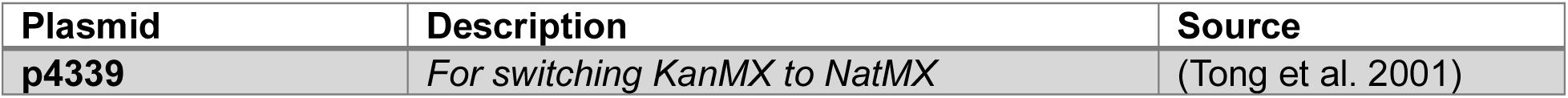
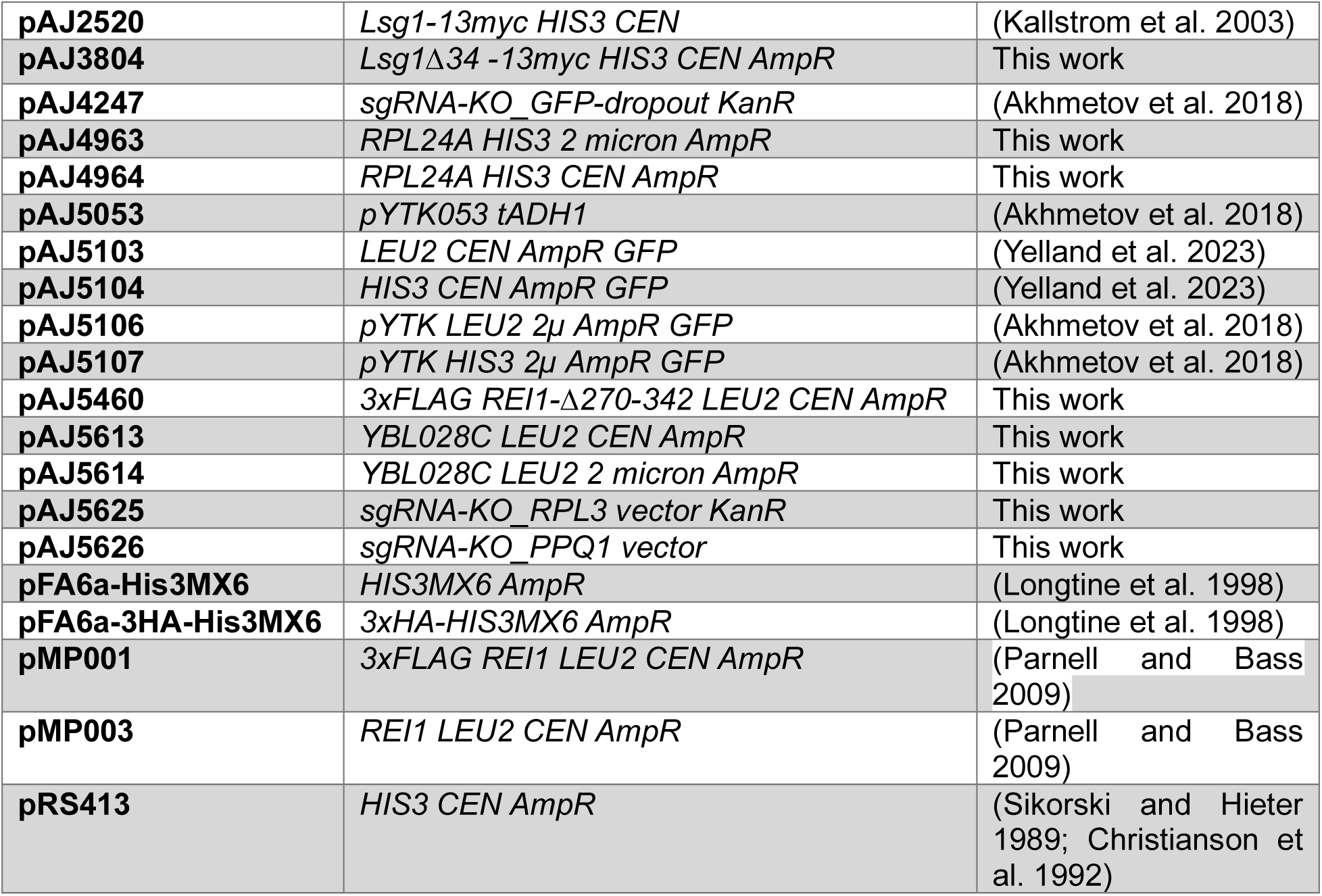
Plasmids used in this work.

All Oligos are purchased from IDT and listed in Table 3.

**Table 3.**
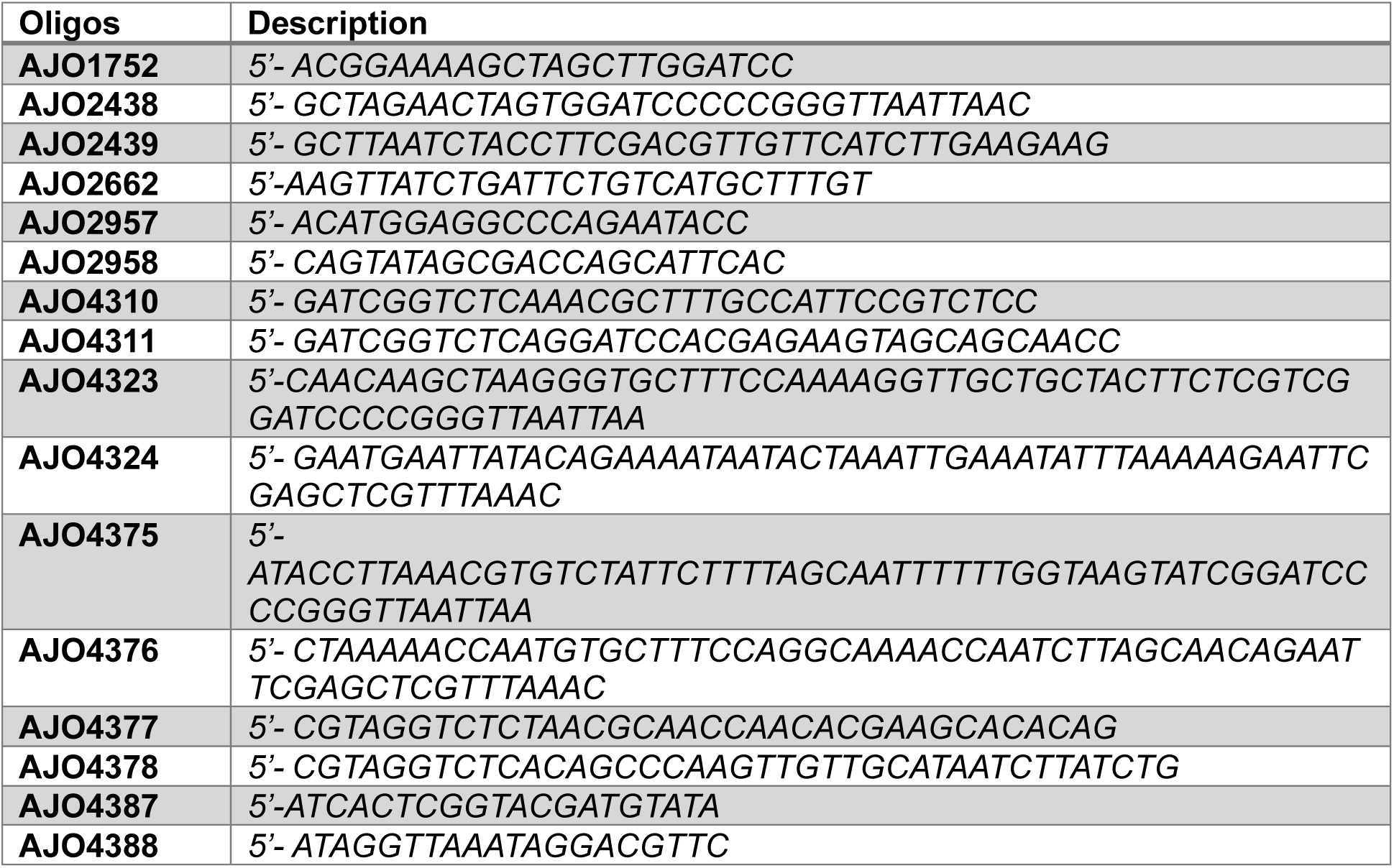
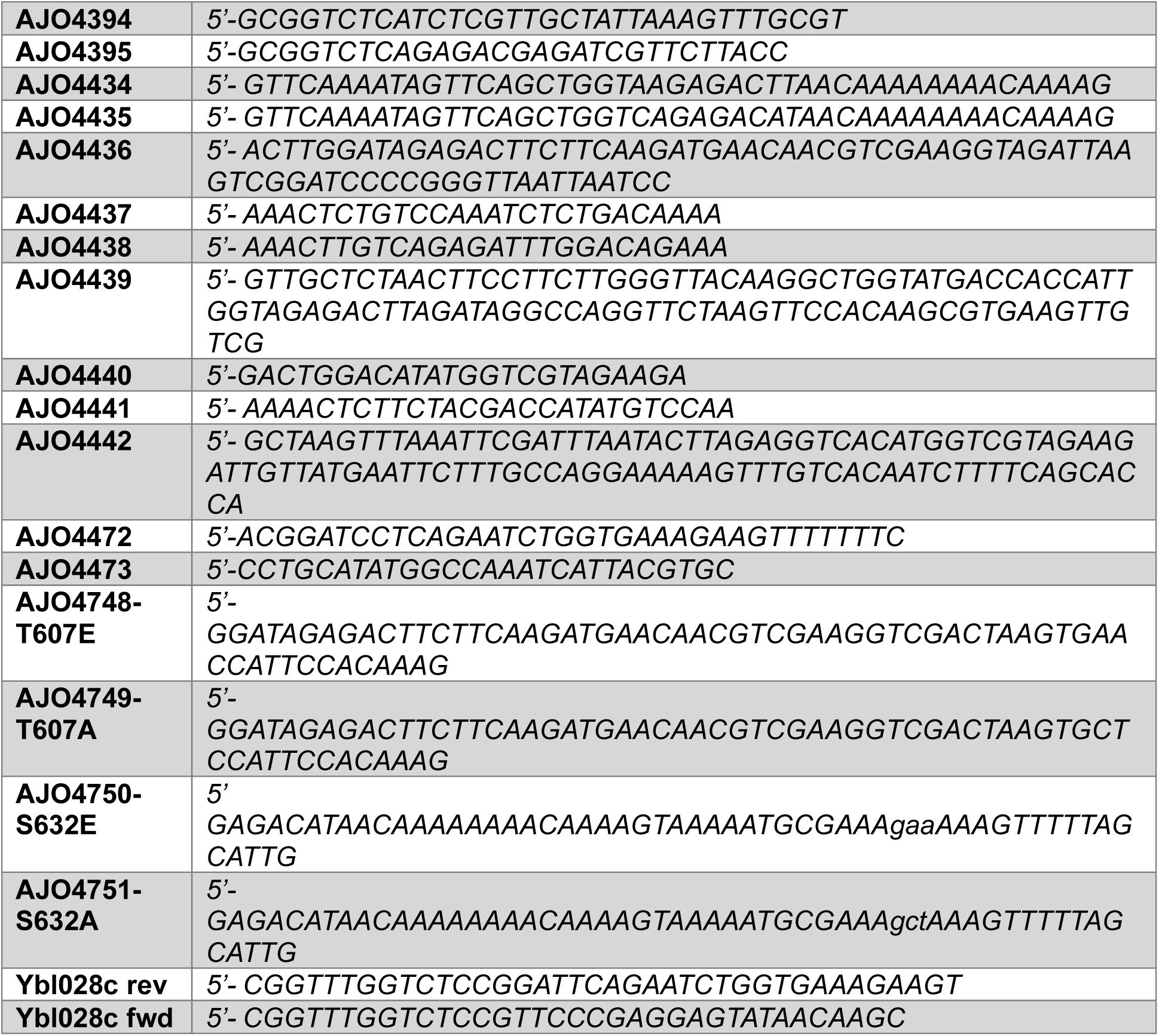
Oligos used in this work.

### Protein Immunoprecipitation for Mass Spectrometry

Independent 250 mL cultures of BY4741 (*WT*), AJY1872(*LSG1-TAP*), and AJY4686 (*LSG1-TAP rei1Δ reh1Δ arx1-S347P*) were grown in triplicate in YPD at 30°C until OD_600_ ∼ 0.4. Cells were harvested by centrifugation (4000 x g for 5 min) and stored at −80°C. The cell pellets were washed with 1 mL ice-cold IP buffer (20 mM Tris-HCl pH 7.5, 100 mM KCl, 10 mM MgCl_2_, 5 mM β-mercaptoethanol (BME), 100 µg/mL cycloheximide(CHX), 1 mM PMSF, 1 µM pepstain, 1 µM leupeptin and EDTA-free Protease Inhibitors Tablets (1 tablet/10mL) (Fisher Scientific)), then resuspended in 400 µL IP buffer. Cells were lysed with 0.5 mm diameter grass beads (Biospec.com) by three cycles of 30s vortexing followed by 1 min on ice. Lysates were separated from the beads by centrifugation and combined with two beads washes (∼1 mL in total). NP40 was added to a final concentration of 0.1% and the lysates were clarified by centrifugation for 10 min at 18,000 x g, 4°C.

Rabbit IgG coupled Dynabeads, prepared in house, were added to equal amounts of extracts based on A_260_ and incubated for 90 min at 4°C with rotation. Beads were washed three times with IP buffer (without NP40). Bound complexes were eluted with TEV protease for 2 hours at 4 °C. Eluates were mixed with Laemmli buffer, briefly electrophoreses in an SDS–PAGE gel, and the bulk protein band digested *in gel* with sequencing-grade trypsin digestion and peptides were prepared and analyzed by LC–MS/MS analysis as previously described (cite Ruta’s current Nat Comms ms). The resulting spectra were processed with Proteome Discoverer version 2.5 (Thermo Scientific) using the Sequest HT search engine, with 10 ppm mass tolerance for the MS at 0.6 Da for the MS/MS. Identifications were validated with Scaffold 5 (Proteome Software). Raw data were processed with MaxQuant and volcano plots were generated using Graphpad Prism software.

### Serial dilution assay

A single freshly grown colony from each strain was resuspended in sterile water to a density of 1×10^7 cells per mL. Ten-fold serial dilutions were prepared in sterile water and 5 microliters per dilution was spotted onto YPD media. Cells were then grown at 30°C for the indicated times.

### Ribosome sedimentation assay

Fresh overnight cultures of yeast strains carrying the desired plasmids were subcultured in 50 mL selective medium to give a starting OD600 of 0.05–0.1. Cultures were grown for ∼7 h until reaching an OD_600_ of 0.4∼0.5. Cells were harvested by centrifugation (4000 × g, 5 min, 4 °C) and stored at –80 °C. Cell pellets were washed with 1 mL ice-cold TMNB buffer (20 mM Tris-HCl pH 7.5, 100 mM NaCl, 10 mM MgCl2, 5 mM β-mercaptoethanol, 1 mM PMSF, 1 µM pepstatin, 1 µM leupeptin, and 1 tablet/10 mL EDTA-free protease inhibitor (Fisher Scientific)) and resuspended in 200 µL TMNB buffer. Cells were lysed with 0.5 mm glass beads (Biospec) by seven cycles of 30 s vortexing followed by 1 min on ice. Lysates were separated from the beads by centrifugation and pooled with 100 µL bead wash (∼300 µL total). Lysates were clarified by centrifugation at 18,000 x g for 10 min at 4 °C.

Clarified extracts were normalized based on A_260_ and 50 uL of extract was layer over a 50 µL sucrose cushion (17% sucrose in TMNB buffer) in a 200 uL ultracentrifuge tube (Beckman Coulter, cat. no. 343775). Samples were centrifuged in a TLA-100 rotor (Beckman Coulter) at 70k rpm for 20 min at 4°C.

After centrifugation, the top 50 µL was transferred to a clean microtube (supernatant fraction). The middle 40 µL was removed and the remaining 10 uL containing the pellet was resuspended in 40 µL TMNB buffer (pellet fraction). 12.5 µL of 5x Laemmli buffer was added and samples were separated on a 6–18% gradient SDS-PAGE gel, followed by western blot analysis. Alternatively, the pellet fraction was submitted directly for mass spectrometry analysis.

### Polysome profiles

Yeast cultures were grown as described above for Ribosome Sedimentation Assay except when cultures reached an OD_600_ of 0.4∼0.5, cycloheximide was added to a final concentration 0.1 mg/mL. Cultures shaken for a further 10 min at 30 °C and then harvested by centrifugation (4000 × g, 5 min, 4 °C). Cells were immediately lysed in TMNB buffer as described above. Clarified extracts were snap frozen and stored at −80°C. Five A_260_ units of clarified extract were loaded onto 10%-50% (w/v) sucrose gradients (containing 50 mM Tris-Acetate pH 7.0, 50 mM NH_4_Cl, 12 mM MgCl_2,_ 1 mM PMSF, 1 µM pepstatin, 1 µM leupeptin, and 1 tablet/10 mL EDTA-free protease inhibitor (Fisher Scientific)) and centrifuged for 75 min at 50,000 rpm in a Beckman SW55 rotor.

Gradients were fractionated using a BioComp Piston Gradient Fractionator fitted with a Triax flow cell into 400 μl fractions with continuous monitoring at 260nm. 40 µL 100% TCA was added to each fraction, mixed and stored at −20°C overnight. Fractions were centrifuged at 4°C for 15min at 18,000 x g and pellets were resuspended in 25 µL 1XSDS-PAGE sample buffer and heated at 99 °C for 3mins. Proteins were separated on 6-18% gradient SDS-PAGE gels, transferred to nitrocellulose membrane, and subjected to western blot analysis using the indicated antibodies as described (western blot analysis).

### Western blot analysis

Primary antibodies used in this study were mouse anti-FLAG (Thermo Fisher Scientific, Dilution 1:10,000), rabbit anti-Rpl8 (K.-Y. Lo, Dilution 1:20,000), mouse anti-HA (Biolegend, Dilution 1:5000) and anti-Lsg1 (from lab). Secondary antibody was Goat anti-Rabbit antibody-IRDye 680LT (Li-Cor Biosciences, Dilution 1:20,000) or Goat anti-Mouse antibody-IRDye 800CW (Li-Cor Biosciences, Dilution 1:15,000). Blots were imaged with an Odyssey CLx infrared imaging system (Li-Cor Biosciences) using Image Studio (Li-Cor Biosciences).

### Spontaneous Suppressor Screen

To identify spontaneous suppressors, twelve independent cultures of AJY3027 were grown to saturation in a 48 well plate in 250 uL YPD medium supplemented with 75 ug/mL ampicillin at room temperature with shaking. Cells were subcultured and grown similarly to saturation three additional times. Final cultures were plated on YPD medium to identify faster growing clones. Single isolates of fast-growing clones from five independent cultures were subjected to whole genome sequencing and SP analysis (Novogene, Sacramento, CA).

## Results and Discussion

### Rpl24 is deficient from 60S in *rei1Δ reh1Δ* cells

Our current understanding of pre-60S maturation in the cytoplasm suggests that Rei1 is recruited to the pre-60S by eL24 which provides a binding site for Rei1 (Lebreton et al. 2006). Rei1 then promotes the release of Arx1 and Alb1 (Figure 1A) (Hung and Johnson 2006; Lebreton et al. 2006). However, Rei1 and its paralog Reh1 appear to have an additional redundant function in 60S biogenesis (Parnell and Bass 2009). To gain insight into the nature of the pre-60S defect in *rei1Δ reh1Δ* mutant cells, we characterized the protein composition of pre-60S subunits from wild-type and *rei1Δ reh1Δ* cells. We chose to affinity purify subunits via Lsg1, which remains bound to subunits during successive maturation steps in the cytoplasm (Kargas et al. 2019; Zhou et al. 2019). We affinity purified Lsg1 from wild-type and *rei1Δ reh1Δ arx1-S347P* mutant cells and identified associated proteins by mass spectrometry (Figure 1B). We included the *arx1-S347P* mutation, which complements loss of *ARX1* but bypasses the requirement for *REI1* (Lo et al. 2010), to avoid issues associated with the role of Arx1 in nuclear export (Bradatsch et al. 2007; Hung et al. 2008).

**Figure 1.**
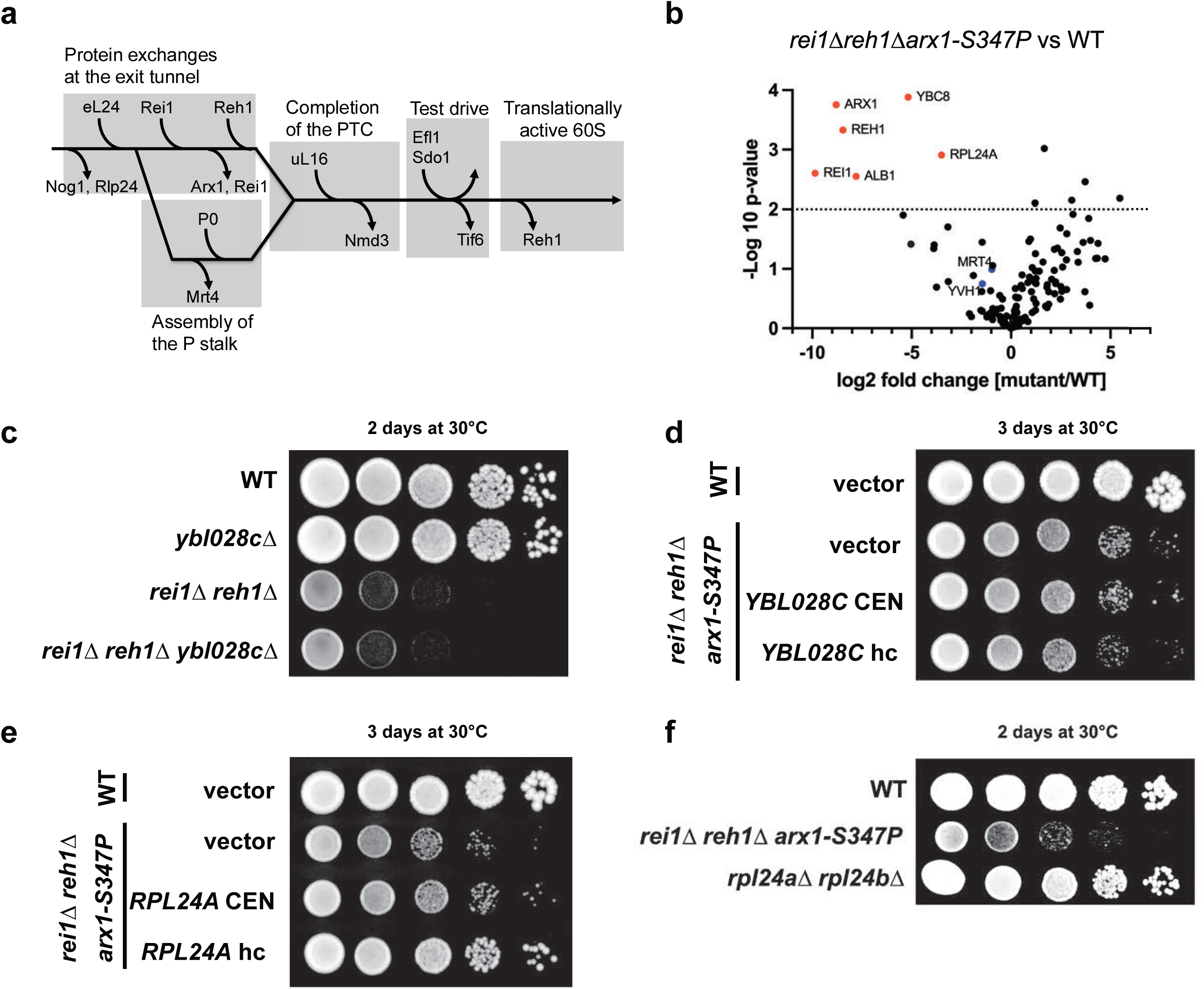
**A)** Schematic representation of the cytoplasmic maturation pathway of the 60S ribosomal subunit. Key maturation events are highlighted, including exchange of factors at the polypeptide exit tunnel, P-stalk assembly, completion of the peptidyl transferase center (PTC), and the "test drive" step that licenses the subunit for translation. **B)** Mass spectrometry analysis of affinity-purified 60S subunits using TAP-tagged Lsg1 from wild-type (*WT*) and *rei1Δ reh1Δ arx1-S347P* mutant strains. **C)** Serial dilution growth assays using *WT*, *ybl028cΔ*, *rei1Δ reh1Δ*, and the triple mutant *rei1Δ reh1Δ ybl028cΔ.* **D)** *rei1Δ reh1Δ arx1-S347P* mutant strains were transformed with an empty vector, a low copy (CEN) plasmid expressing *YBL028C* (pAJ5613), or a high-copy (hc) plasmid expressing *YBL028C* (pAJ5614). **E)** Serial dilutions of *WT* and *rei1Δ reh1Δ arx1-S347P* strains transformed with an empty vector, a CEN plasmid expressing *RPL24A* (pAJ4964), or a high-copy plasmid expressing *RPL24A* (pAJ4963). **F)** Serial dilutions of *WT*, *rei1Δ reh1Δ*, and the double deletion mutant *rpl24aΔ rpl24bΔ*.

Mass spectrometry of Lsg1-bound pre-60S from *reh1Δ rei1Δ arx1-S347P* cells gave a protein profile very similar to that from wild-type cells, but with some notable differences (Figure 1B and Supplementary Table 1). As expected, Reh1 and Rei1 were strongly depleted, reflecting their genetic absence. Arx1 and its associated protein Alb1 (Lebreton et al. 2006) were also strongly depleted. Although Rei1 is normally needed for the release of Arx1, the *arx1-S347P* mutation bypasses the requirement for Rei1. Thus, the depletion of Arx1 likely reflects its spontaneous release from pre-60S after it enters the cytoplasm. In addition, Ybl028c, a protein of unknown function but which has been observed in previous pre-60S structures (Kater et al. 2020; Zhang et al. 2023), was strongly depleted. Lastly, the ribosomal protein eL24 was also strongly depleted. In contrast, the levels of the assembly factors Yvh1, Mrt4, Tif6 and Nmd3 were not significantly affected (Supplementary Table 1), suggesting that their release from the pre-60S was independent of the defect of *rei1Δ reh1Δ* cells.

Considering the depletion of Ybl028c and eL24, we tested for genetic interaction between *YBL028C* and *REI1 and REH1*. No obvious growth defect was observed for the *ybl028cΔ* strain and no synthetic genetic interaction was seen between the deletion of *YBL028C* and the deletion of *REI1* and *REH1* (Figure 1C). Furthermore, high-copy expression of *YBL028C* did not affect the growth of *rei1Δ reh1Δ* cells (Figure 1D).

### eL24 levels are restored by high-copy *RPL24A*

We next focused on eL24. The depletion of eL24 was unexpected because eL24 is reported to recruit Rei1 to the pre-60S (Lebreton et al. 2006). In that model, eL24 should load onto pre-60S before and independently of Rei1. Because we observed loss of eL24 from pre-60S in the absence of Rei1 and Reh1, we determined if high-copy expression of eL24 could suppress the growth defect of *rei1Δ reh1Δ* cells. In yeast, eL24 is expressed from two paralogs, *RPL24A* and *RPL24B* (Baronas-Lowell and Warner 1990), which differ by only 5 out of 155 residues. We cloned *RPL24A* (which is reported to have higher expression than *RPL24B* (Baronas-Lowell and Warner 1990) into a high-copy vector. We found that overexpression of *RPL24A* from a high-copy vector, but not a low copy centromeric vector, partially suppressed the growth defect of *rei1Δ reh1Δ* cells (Figure 1E). The depletion of eL24 from subunits in the absence of Rei1 and Reh1 and suppression by high-copy *RPL24A* suggested that the loading of eL24 is impaired in these cells.

To test if overexpression of *RPL24A* actually led to increased levels of eL24 in 60S subunits, we sedimented extracts from WT cells containing empty vector and from *rei1Δ reh1Δ arx1-S347P* cells containing empty vector or high-copy *RPL24A*, through sucrose cushions. We then applied mass spectrometry to compare the relative amounts of eL24 on ribosomes from the three strains. We normalized the values for eL24 relative to 60S subunit proteins, using the top 10 proteins with the highest spectral counts, from WT cells, which we assumed were fully loaded with eL24 (Supplementary Table 2). We recapitulated the result that eL24 was underloaded in *rei1Δ reh1Δ* cells (Figure 2A). Importantly, we found that eL24 levels were increased when *RPL24A* was overexpressed (Figure 2A). Although the eL24 deficit in this experiment was less than that observed in Figures 1 and 3, the restoration of eL24 levels by high copy *RPL24A* is consistent with our interpretation that Rei1 and Reh1 promote the loading of eL24.

**Figure 2.**
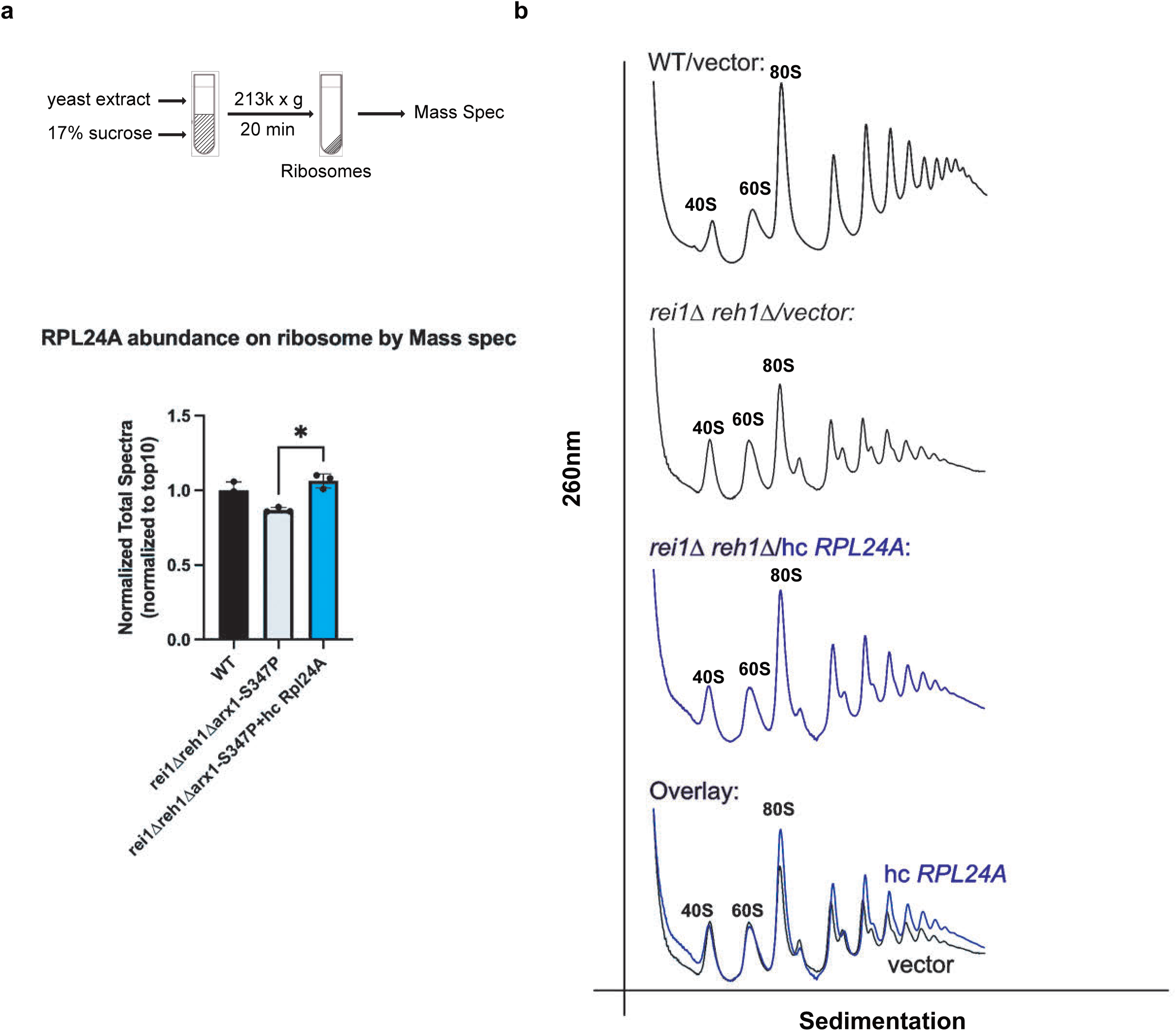
High-copy *RPL24A* partially restored the eL24 levels and translation defect. **A**) Semiquantitative mass spectrometry of ribosomes from *WT* with empty vector or *rei1Δ reh1Δ arx1-S347P* strains transformed with either an empty vector or high-copy (hc) *RPL24A*. **B)** Polysome assay of *WT,* or *rei1Δ reh1Δ arx1-S347P* strains transformed with either an empty vector (black) or a high-copy (hc) *RPL24A* plasmid (blue). The bottom panel shows the overlay of polysome profiles comparing *rei1Δ reh1Δ arx1-S347P* strains carrying an empty vector versus the high-copy (hc) *RPL24A* plasmid.

**Figure 3.**
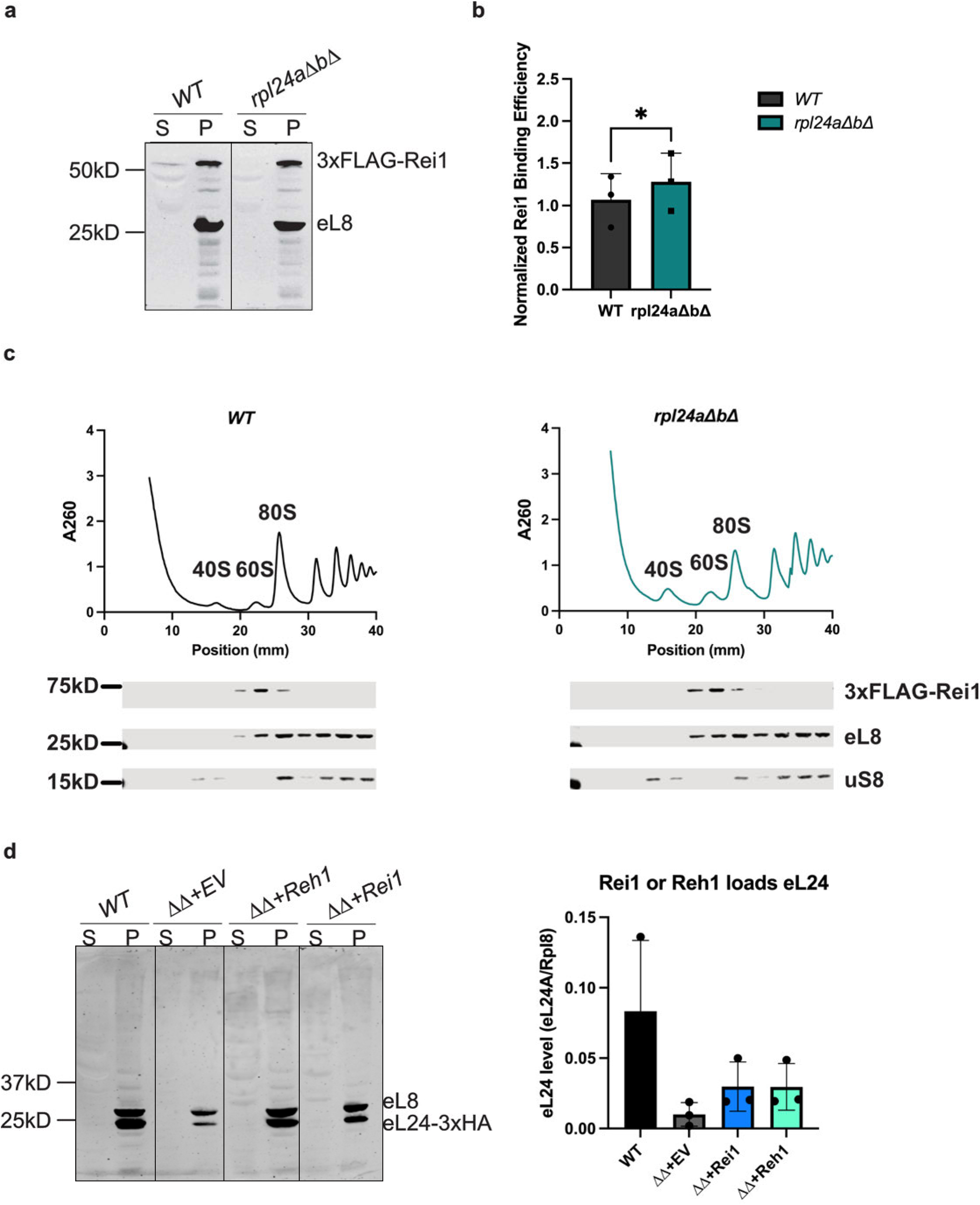
**A)** Western blot analysis of yeast extracts from *WT* strain (BY4741) or r*pl24aΔ rpl24bΔ* (*rpl24aΔbΔ*) strain (AJY1544) containing 3XFLAG-Rei1, following a ribosome sedimentation assay. Supernatant (S) or pellet (P) was collected and probed with anti-FLAG (for Rei1) and anti-eL8 antibodies. **B)** Quantification of Rei1 level in *WT* or *rpl24aΔbΔ* strains from (A). Rei1 signal relative to eL8 was normalized to the Rei1 vs eL8 in *WT*. **C)** Confirmation that 3xFLAG-Rei1 co-sediments with 60S subunits. Polysome profiling was performed on extracts from *WT* (black) and *rpl24aΔbΔ* (teal) strains, followed by Western blot analysis of fractions using antibodies against FLAG (Rei1), eL8 (60S marker), and uS8 (40S marker). **D)** Rei1 and Reh1 can each facilitate eL24 loading onto the ribosome. WT: *RPL24A*-3xHA tagged strain (AJY4809) with empty vector, ΔΔ: *rei1Δ reh1Δ arx1-S347P RPL24A-*3xHA-tagged strain (AJY4810) carrying empty vector (+EV), a vector expressing Reh1 (+Reh1) or Rei1 (+Rei1). Yeast extracts were subjected to sucrose cushion sedimentation followed by western blotting to assess eL24 levels.

We next determined if high-copy *RPL24A* restored the translation defect of *rei1Δ reh1Δ arx1-S347P* cells. Extracts were prepared from *rei1Δ reh1Δ arx1-S347P* cells containing either empty vector or high-copy *RPL24A*. Cells lacking Rei1 and Reh1 displayed elevated levels of free 60S and 40S subunits relative to monosomes (Figure 2B), indicative of a subunit joining defect, as described previously (Parnell and Bass 2009). These cells also contained significant populations of half-mers, representing mRNAs containing unjoined 40S subunits, and overall reduced levels of polysomes. The presence of high-copy *RPL24A* resulted in a modest increase in monosomes relative to free subunits, indicating more efficient utilization of the subunits, a modest reduction in half-mers and a slight increase in polysomes (Figure 2B).

### The reported eL24 binding site on Rei1 is not required for eL24 loading

We sought to test whether the recruitment of eL24 to pre-60S required its physical interaction with Rei1. An interaction between eL24 (Rpl24B) and amino acids 270-342 of Rei1 was reported previously, using a yeast 2-hybrid assay (Lebreton et al. 2006). However, we were unable to recapitulate this result, and the original vectors and their sequences are no longer available. It is possible that subtle differences in constructs or strains account for our inability to detect an interaction by yeast 2-hybrid. We proceeded to test if deleting the reported interaction domain on Rei1 would inhibit eL24 recruitment. *rei1Δ*270-342 was unable to complement the growth defect of *rei1Δ reh1Δ* mutant cells (Figure S1B). However, *rei1Δ270-342* did promote the assembly of eL24 into 60S subunits (Figure S1C, D). Thus, the putative binding site for eL24 on Rei1 is not required for eL24 loading into ribosomes. The lack of complementation was unexpected and may be related to the functional interaction between Rei1 and Arx1, as the region that was deleted includes the Arx1 interaction helix (Greber et al. 2016).

### Deletion of eL24 does not impair Rei1 loading onto the ribosome

The deficit of eL24 in pre-60S particles and the ability of high-copy *RPL24A* to suppress *rei1Δ reh1Δ arx1-S347P* led us to consider the possibility that Rei1 and Reh1 promote the loading of eL24, a model that is contrary to the prevailing understanding that eL24 recruits Rei1. If Rei1 recruits eL24, the binding of Rei1 to pre-60S should be independent of eL24. We revisited our previous observation that Rei1 could bind 60S in the absence of eL24 (Lo et al. 2010). Using a sucrose cushion sedimentation assay, we found that Rei1 pelleted with ribosomes in the absence of eL24 (Figure 3A). Indeed, the abundance of Rei1 in the pellet fraction from the *rpl24aΔbΔ* mutant was equal to, if not greater than, that in wild-type cells (Figure 3B). To rule out the possibility that Rei1 was aggregating and sedimenting independently of 60S, we monitored the sedimentation of Rei1 in sucrose density gradients from *WT* and *rpl24aΔbΔ* mutant cells (Figure 3C). In both *WT* and *rpl24aΔbΔ* mutant cells, Rei1 cosedimented exclusively with free 60S subunits. These results demonstrate that Rei1 can bind to pre-60S independently of eL24.

To directly test whether the redundant functions of Rei1 and Reh1 involve eL24, we complemented the *rei1Δ reh1Δ arx1-S347P* strain with either *REI1* or *REH1* expressed in the *RPL24A*-HA background. We then performed sucrose cushion sedimentation followed by western blotting to assess eL24 levels across the four conditions: *WT, REI1* only, *REH1* only, and *rei1Δ reh1Δ arx1-S347P*. As shown in Figure 3D (left), eL24 levels were significantly increased and restored to comparable levels upon complementation with either Rei1 or Reh1, although not fully to wild-type levels. This result strongly suggests that the redundant function of Rei1 and Reh1 is the loading of eL24.

### Extragenic suppressors of *rei1Δ reh1Δ*

So far, we have provided evidence that Rei1 and Reh1 facilitate the loading of eL24. However, if that was their sole redundant function, we would expect that the growth of the *rei1Δ reh1Δ* double mutant would be equivalent to that of a *rpl24aΔ rpl24bΔ* double mutant. This is not the case; the *rei1Δ reh1Δ* double mutant has a much greater growth defect (Figure 1F), suggesting that these proteins have yet an additional role in ribosome assembly. To identify this additional function of Rei1 and Reh1, we screened for bypass suppressors of the slow-growth defect of *rei1Δ reh1Δ* double mutants. We did this in the absence of *ARX1* to avoid isolating mutations in *ARX1* that suppress *rei1Δ* (Hung and Johnson 2006). Independent cultures of strain AJY3027 deleted of *REI1*, *REH1* and *ARX1* were serially subcultured multiple times and then plated to identify faster growing clones. Whole genome sequencing of five independently derived fast-growing clones identified two independent mutations in *LSG1*, and single mutations in *RPL3* and *PPQ1* (Table 4). We were unable to identify the suppressing mutation in one clone.

**Table 4.** Spontaneous suppressors of *rei1Δ reh1Δ arx1Δ*.

| Gene Name | Nucleotide Position | Nucleotide Change | Amino Acid Position | Amino Acid Change |
| --- | --- | --- | --- | --- |
| *LSG1* | 1862 | A>T | 621 | H>L |
| *LSG1* | 1855 | A>C | 619 | K>Q |
| *RPL3* | 170 | T>G | 57 | V>G |
| *PPQ1* | 1471 | G>T | 491 | G>C |

Lsg1 is a cytoplasmic GTPase that associates with nascent pre-60S in the cytoplasm, where it is required for the release of the nuclear export adapter protein Nmd3 (Hedges et al. 2005). Although we have determined the structure of the G-domain of Lsg1 in complex with 60S (Malyutin et al. 2017; Zhou et al. 2019), the two mutations we identified in Lsg1 (K619Q and H621L) map to the extreme C-terminus which has not been resolved in structures. The ribosomal protein uL3 (encoded by *RPL3*) assembles into the pre-60S in the nucleus. Intriguingly, the mutated residue V57 lies in a cleft within uL3 that embraces the N-terminus of eL24 (Figure S2). Ppq1 is a cytoplasmic protein phosphatase but its substrate(s) is not known. Ppq1 has been reported to impact translation but has not been functionally associated with ribosome assembly (Chen et al. 1993). The suppressing mutation, G491C, affects a residue in the vicinity of the active site suggesting that it might compromise the phosphatase activity of Ppq1.

To confirm that the mutations identified are sufficient for suppression of *rei1Δ reh1Δ*, we reconstructed these mutations *de novo* in a strain deleted of *REI1* and *REH1* and carrying the *arx1-S347P* mutation, ruling out the possibility that the suppression was specific for the deletion of *ARX1*. All reconstructed mutants showed faster growth than the parental mutant strain, demonstrating that the identified mutations were causative (Figure 4A).

**Figure 4.**
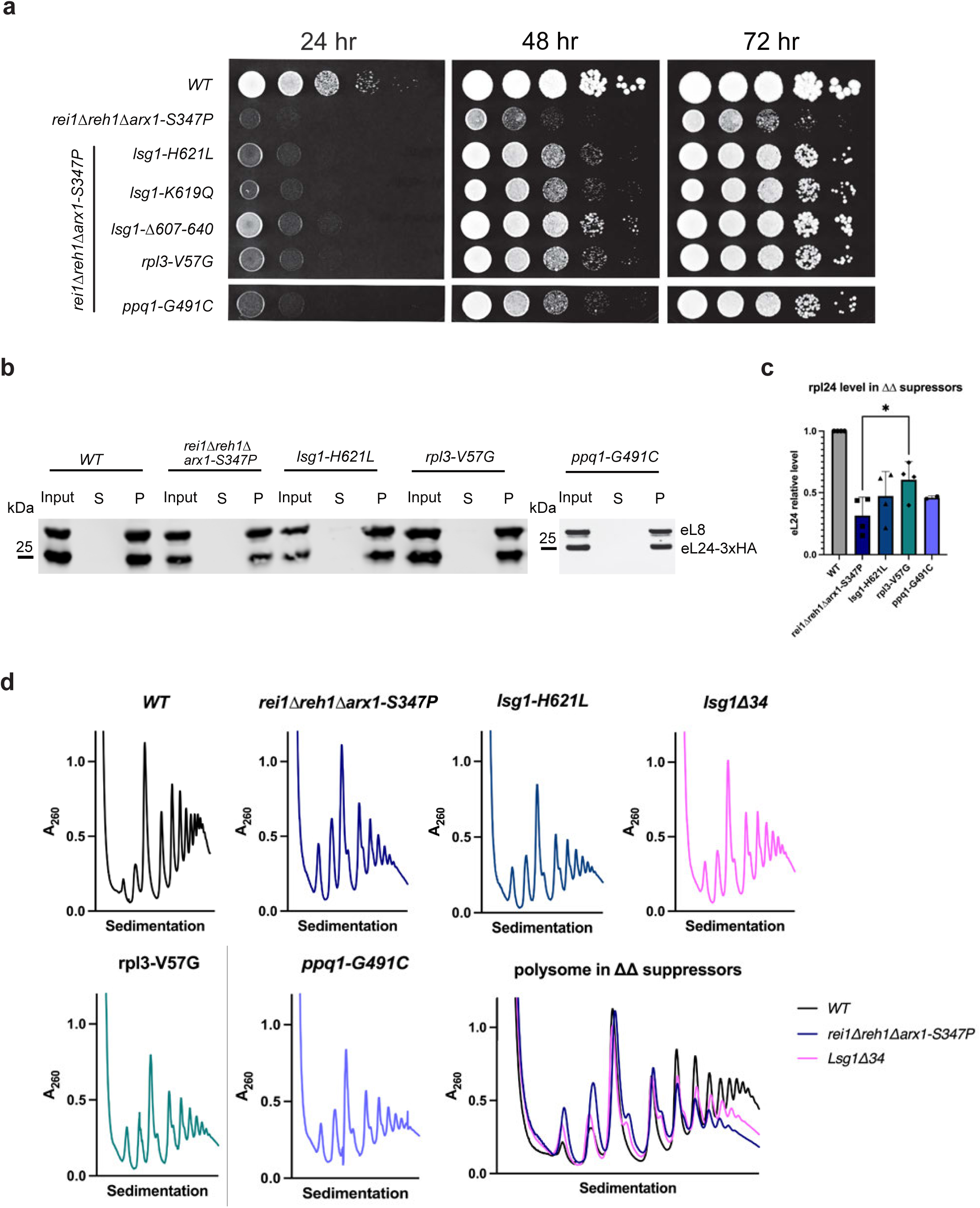
**A)** Serial dilution assays were performed to confirm the growth suppression phenotypes of the mutants in the *rei1Δ reh1Δ arx1-S347P* background. **B)** Yeast extracts from WT strain or *rei1Δ reh1Δ arx1-S347P* or suppressor strains were subjected to a sucrose cushion assay. Supernatant (S) and pellet (P) fractions were analyzed by western blot using anti-HA (for HA-tagged eL24) and anti-Rpl8 antibodies. **C)** Quantification of eL24-HA levels in WT and suppressor strains. HA signal was normalized to Rpl8, indicating that eL24 loading onto ribosomes was partially restored in all suppressor mutants. **D)** Polysome assays were performed across WT strain or *rei1Δ reh1Δ arx1-S347P* or suppressor strains. Subtle restoration of polysome levels was observed in the suppressor backgrounds.

We also determined if eL24 levels were restored in a subset of the suppressor mutants. To this end, we integrated a 3xhemagglutinin (3xHA) epitope tag at the C-terminus of the genomic *RPL24A* locus in *WT*, *rei1Δ reh1Δ arx1-S347P* cells as well as the *rei1Δ reh1Δ arx1-S347P* double mutant suppressed by *lsg1-H621L*, *rpl3-V57G* or *ppq1-G491C*. We then sedimented cell extracts through sucrose cushions as described above and analyzed the input and pellet fractions for eL24-HA, normalizing the values to eL8 (Figure 4B). In all cases we observed partial restoration of eL24 levels in subunits in the suppressed strains (Figure 4C), indicating that the suppressing mutations enhance eL24 loading in the *rei1Δ reh1Δ* mutant. In addition, the suppressors partially restored the translation defect of *rei1Δ reh1Δ* cells, as shown by the modest decrease in free subunit levels as well as a slight increase in polysomes.

### Deletion of the C-terminus of Lsg1 suppresses the growth defect of *rei1Δ reh1Δ* mutant cells

The C-terminus of Lsg1 is highly basic and also highly conserved throughout eukaryotes, with the mutated residue H621 being nearly invariant in eukaryotes (Figure 5A) The positive charge of the C-terminus suggests that it might be involved in ribosome binding through interaction with RNA. Because both suppressing mutations *lsg1-H621L* and *lsg1-K619Q* affected the C-terminus of Lsg1, and could affect ribosome binding, we tested if deleting the C-terminus of Lsg1 would also suppress *rei1Δ reh1Δ arx1-S347P* cells. Deletion of the C-terminal 34 amino acids (aa607-640) of Lsg1 (*lsg1Δ34*) did not cause an obvious growth defect on its own (Figure 5B). However, *lsg1Δ34* suppressed the growth defect of *rei1Δ reh1Δ arx1-S347P* cells when integrated into the genome or expressed from a low copy vector (Figure 4A, 5C). In fact, deleting the C-terminal 34 amino acids resulted in stronger suppression than that observed for any of the spontaneous point mutants that we identified in our screen (Figure 4D). To test if Lsg1Δ34 also showed reduced binding to ribosomes, we sedimented extracts through sucrose cushions and monitored the presence of Lsg1 in the soluble (free) vs pellet (ribosome-bound) fractions. We found that whereas approximately 80 percent of wild-type Lsg1 was ribosome-bound, this decreased to approximately 60 percent for Lsg1Δ34, indicating that the truncated protein had reduced affinity for pre-60S (Figure 5D).

**Figure 5.**
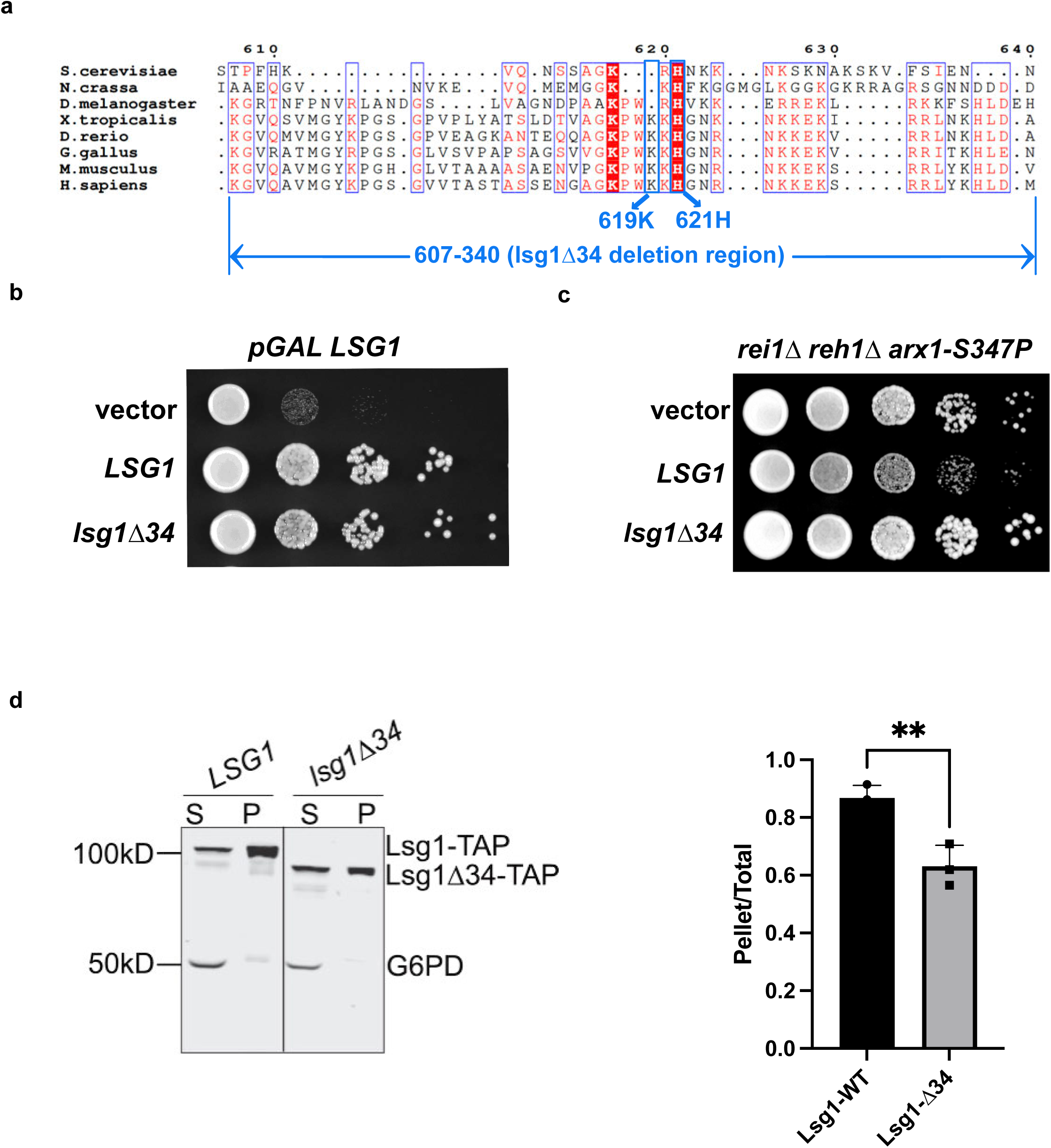
**A)** Multi-sequence alignment of Lsg1 and its homologs. Sequences were aligned using T-coffee, then output using ESPript 3.0. Aligned sequences: protein, organism (access No.): Lsg1, *S. cerevisiae S288C* (NP_011416.3); Large subunit GTPase 1 homolog, *N.crassa* (KAK3952141.1); Large subunit GTPase 1 homolog, *D. melanogaster* (Q9W590.1); large subunit GTPase 1 homolog, *X. tropicalis* (XP_002937880.1); Lsg1 *D.rerio*, (NP_997807.1); large subunit GTPase 1 homolog, *G. gallus*, (Q5ZJD3.1); large subunit GTPase 1 homolog isoform 1, *M. musculus* (NP_835170.1); large subunit GTPase 1 homolog, *H. sapiens* (NP_060855.2). **B)** Serial dilutions of *pGAL LSG1* (AJY3827) containing empty vector, or vectors expressing *LSG1* WT or *lsg1*Δ34 were plated on SC His- medium containing glucose and incubated for 2 days at 30°C. **C)** Empty vector or vectors expressing *LSG1* or *lsg1Δ34* were transformed into the *rei1Δ reh1Δ arx1-S347P* strain (AJY4686) and serial dilutions were plated on SC His-medium containing glucose and grown for 2 days at 30°C. **D)** Left: Sucrose cushion sedimentation assay for Lsg1-ribosome association. *WT* (BY4741) transformed with *LSG1*-WT or *lsg1Δ34* vector. A representative western blot of supernatant (S) and pellet (P) fractions from extracts of WT cells expressing Lsg1 or Lsg1Δ34. Right: Quantification of fraction of that was ribosome-bound. The amount of ribosome-associated Lsg1 was normalized to the WT level (set to 1), indicating that Lsg1 loading onto ribosomes was reduced with deletion of the 34 residues at the C-terminus.

The structure of the C-terminus of Lsg1 has not been resolved in cryo-EM studies. However, lysines at positions K619 and K633 are reported to crosslink to lysine 67 of Rei1 (Kargas et al. 2019). Because the sequences of the N-termini of Rei1 and Reh1 are highly similar, it is reasonable to assume that a similar juxtaposition of Lsg1 and Reh1 occurs. AlphaFold3 predicts that residues 587 to 596 of Lsg1, which are highly conserved, form a short alpha helix that passes underneath Reh1 (Figure S3). In low pass-filtered cryo-EM maps of pre-60S particles containing Reh1 (Chitale et al. 2025), we have observed density for an unassigned alpha helix in this position that we ascribe to Lsg1, based on AlphaFold3 modeling. This position would place the C-terminus of Lsg1 in the immediate vicinity of eL24, consistent with the crosslinking data, and its positioning would be constrained by Rei1 and Reh1. We suggest that Rei1 or Reh1 help position the C-terminus of Lsg1 to promote efficient eL24 incorporation.

### Phosphorylation of Lsg1 may influence its interaction with the pre-60S

We next focused on *PPQ1*, in which we identified the suppressor mutation G491C. Ppq1 is a serine/threonine phosphatase related to PP1. The protein contains two distinct domains: a carboxy-terminal phosphatase domain, which shares ∼60% identity with PP1, and an amino-terminal region enriched in serine and asparagine residues. Structural prediction with AlphaFold and alignment with PP1 (PDB: 4MOV) (Figure S4A) show that the Ppq1 C-terminal region (residues 238–549, pink) aligns well with PP1 (cyan). PP1 uses a Mn²⁺ ion for catalysis, and the catalytic center lies adjacent to this Mn²⁺ ion. Although G491 in PPQ1 is not located directly in the catalytic center, it is on the same loop as H485 which coordinates an essential Mn^2+^ ion catalytic residue (Figure S4B). Substitution of glycine with cysteine at position 491 may alter the geometry of the catalytic center (Figure S4B, yellow).

To test whether the suppressing G491C mutation *in PPQ1* may be a loss-of-function allele, we generated a *ppq1Δ* strain. Deletion of PPQ1 did not show an obvious growth defect under standard growth conditions (Figure 6A). We then tested whether *ppq1Δ* could suppress *rei1Δ reh1Δ*. Indeed, *ppq1Δ* suppressed the severe growth defect of *rei1Δ reh1Δ* (Figure 6A), supporting the idea that Ppq1-G491C is a partial loss-of-function mutant.

**Figure 6.**
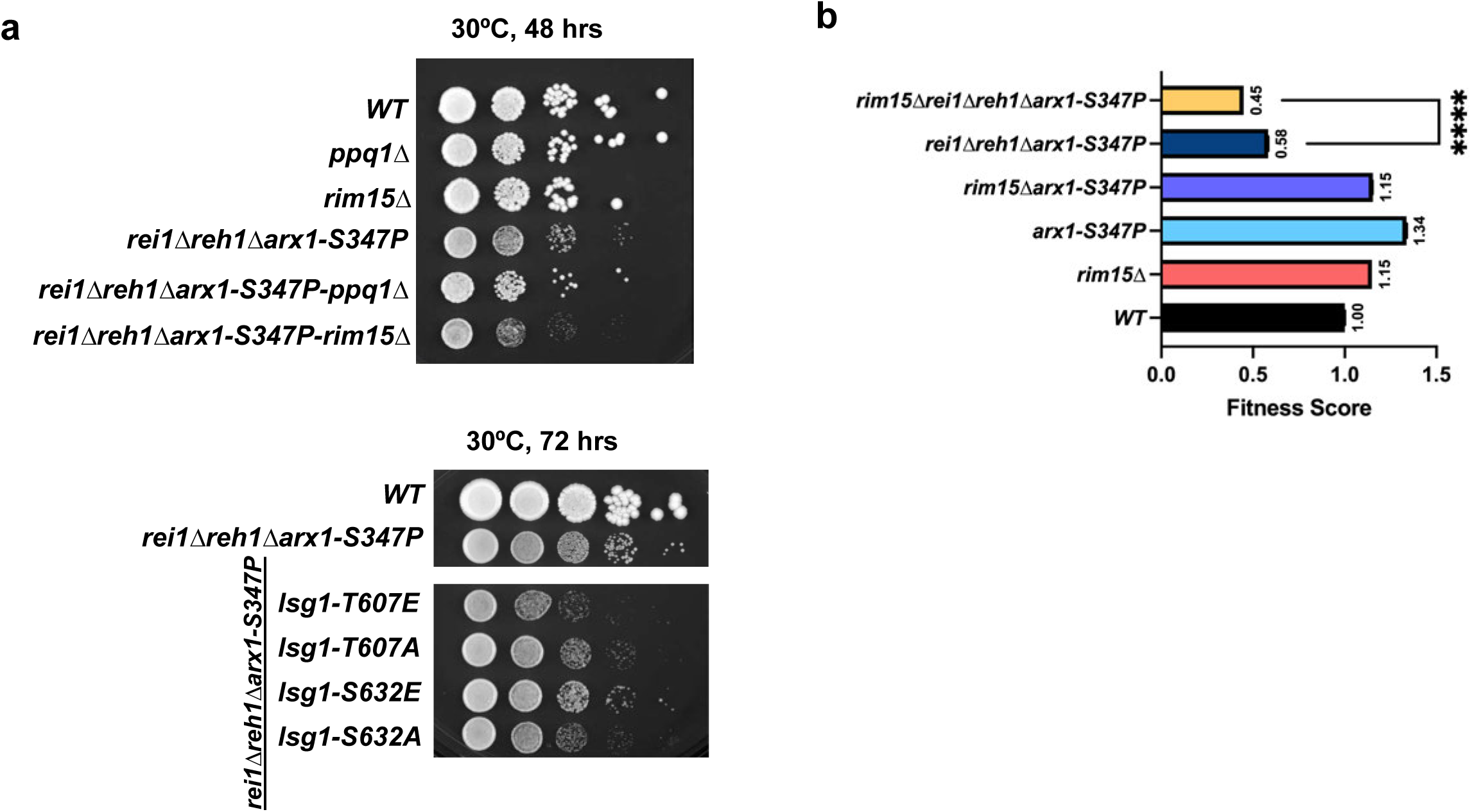
**A)** Serial dilutions of *WT* (BY4741), *ppq1Δ* (AJY4854), *rim15Δ* (AJY4859), *rei1Δ reh1Δ arx1-S347P* (AJY4686), *ppq1Δ rei1Δ reh1Δ arx1-S347P* (AJY4856), and *rim15Δ rei1Δ reh1Δ arx1-S347P* (AJY4860). Cells were spotted on a YPD plate grown for 48 hours at 30°C. Deletion of *PPQ1* suppressed the growth defect of the *rei1Δ reh1Δ arx1-S347P* strain, while deletion of *RIM15* exacerbated it, indicating a negative genetic interaction. **B)** Fitness Score was calculated based on maxim growth rate of each strain in liquid YPD medium at 30°C, WT was set as 1. Statistical significance was assessed by one-way ANOVA followed by Welch’s t test (**** p < 0.0001). WT doubling time was 147.7 min. **C)** Serial dilution spot assays of the indicated strains grown on YPD at 30 °C and imaged after 72 hours. Strains include WT and the *rei1Δ reh1Δ arx1-S347P* background carrying the indicated mutations (*lsg1-T607E*, *lsg1-T607A*, *lsg1-S632E*, *lsg1-S632A*). Cells were serially diluted 10-fold and spotted onto YPD plates. Lsg1 phospho-site mutants did not suppress the growth defect of the *rei1Δreh1Δarx1-S347P* mutant under these conditions.

Although the G491C mutation is likely to affect Ppq1 catalytic activity, no physiological substrates of Ppq1 are known. One candidate is Lsg1, which is reported to be phosphorylated at residues T607 and S632 within the C-terminal 34 amino acids (Holt et al. 2009) and at other residues as well. Phosphorylation of the C-terminus may reduce Lsg1’s affinity for the ribosome, acting similarly to a C-terminal deletion mutant and thereby suppressing the *rei1Δ reh1Δ* phenotype.

To test whether Lsg1 phosphorylation could contribute to its genetic interaction with *rei1Δ* and *reh1Δ* mutations, we introduced phosphomimetic (T607E, S632E) and phosphonull (T607A, S632A) mutations into Lsg1. We also generated a deletion of *RIM15*, the kinase reported to modify T607 (Dokládal et al. 2021). Deletion of *RIM15* resulted in a modest negative genetic interaction with *rei1Δ reh1Δ*. (Figure 6 A, B). The negative genetic interaction with *rim15Δ* is consistent with phosphorylation by Rim15 contributing to the release of the C-terminus of Lsg1. However, in the *rei1Δ reh1Δ arx1-S347P* background, neither of the phosphomimetic mutations suppressed growth, as would have been expected if phosphorylation of these residues promoted release of the C-terminus of Lsg1. Indeed, all of the phosphomimetic and phosphonull mutations resulted in poorer growth in *rei1Δ reh1Δ arx1-S347P* cells (Figure 6D). The modulation of Lsg1 function by phosphorylation is likely to be more nuanced than our analysis of single residues can reveal.

## Discussion

Cytoplasmic maturation of the eukaryotic large ribosomal subunit occurs through an ordered pathway. During this process, the final rproteins are incorporated and assembly factors bind and dissociate in a coordinated manner to promote subunit assembly. Cytoplasmic maturation is initiated by a series of protein exchanges at the exit tunnel, driven by Drg1 which removes Rlp24 and Nog1 (Pertschy et al. 2007; Kappel et al. 2012). The release of Rlp24 makes the eL24 binding site available. It had been proposed that eL24 then binds and recruits the zinc-finger protein Rei1 in yeast (Saveanu et al. 2003). An important function of Rei1 is to recruit the Hsp70 Ssa and its associated J-protein Jjj1 to facilitate the release of Arx1 (Hung and Johnson 2006; Lebreton et al. 2006; Demoinet et al. 2007; Meyer et al. 2007, 2010), a degenerate methionyl aminopeptidase that sits at the exit tunnel and acts as an export factor in yeast (Hung and Johnson 2006; Bradatsch et al. 2007; Hung et al. 2008; Meyer et al. 2010; Bradatsch et al. 2012). Here, we have provided evidence that suggests an alternate order of events. We show that Rei1 recruitment does not depend on eL24 loading. On the contrary, we find that Rei1 is required for the efficient recruitment of eL24.

Most eukaryotes have a single homolog of Rei1, which is ZNF622 in humans. However, in *S. cerevisiae*, Rei1 has a paralog, Reh1, which has functional redundancy with Rei1 (Parnell and Bass 2009). Although Rei1 is required for the efficient release of Arx1, this is a function specific to Rei1 and is not the redundant function of these proteins. Here, we show that Rei1 and Reh1 together are required for the efficient loading of eL24. This conclusion is supported by multiple lines of evidence: eL24 is underloaded in yeast lacking Rei1 and Reh1; high copy expression of *RPL24A*, coding for eL24, suppresses the growth defect of *rei1Δ reh1Δ* cells; and high copy expression of *RPL24A* increases the loading of eL24. However, Rei1 and Reh1 must have a function beyond the loading of eL24 because the yeast deleted of *REI1* and *REH1* have a much greater growth defect than that of *rpl24aΔrpl24bΔ* mutant cells lacking eL24.

To probe the function of Rei1 and Reh1 more deeply, we carried out a screen for spontaneous suppressors of the growth defect of the *rei1Δ reh1Δ* double mutant. We identified mutations in *LSG1*, encoding the cytoplasmic assembly factor Lsg1, as well as in the rprotein gene *RPL3*. Notably, all suppressors partially restored the levels of eL24 on ribosomes, suggesting functions related to eL24 loading. The suppressing mutations in *LSG1* map to the extreme C-terminus of the protein which has not been resolved in published cryo-EM structures. Nevertheless, the structural model suggests that the C-terminus passes beneath Rei1 and Reh1, consistent with previously published protein crosslinking data(Sailer et al. 2022). The mutation in Rpl3 lies in a cleft that accepts the N-terminus of eL24 and is close in space to the predicted position of the C-terminus of Lsg1. We propose that the interaction of Lsg1 with Reh1 and Rei1 positions the C-terminus of Lsg1 to promote the loading of eL24. Because the deletion of the C-terminus of Lsg1 also suppresses the growth defect of *rei1Δ reh1Δ* mutant cells and reduces the affinity of Lsg1 for 60S, Rei1 and Reh1 may also modulate the release of Lsg1. The retention of Lsg1 on subunits could contribute to the strong growth defect of these cells.

The identification of a suppressing mutation in *PPQ1* was unexpected. Ppq1 is a PP1 family protein phosphatase, but its substrate is unknown. It has not previously been linked to ribosome assembly, although mutations in PPQ1 (SLA6) were initially identified as allosuppressors which enhanced the suppression of translational suppressor mutations (Song and Liebman 1987; Vincent and Liebman 1994), suggesting that Ppq1 impacts translation in some manner. Finding a loss of function mutation in *PPQ1* as a suppressor of *rei1Δ reh1Δ* strongly suggests a role for Ppq1 in assembly. Lsg1 is reported to be phosphorylated and is a potential substrate for Ppq1. In particular, the phosphorylated residues T607 and S632 are in the extreme C-terminus, deletion of which weakens the association of Lsg1 with 60S and suppresses *rei1Δ reh1Δ*. It is possible that phosphorylation of the C-terminus modulates its binding to the 60S. In this case, loss of Ppq1 function would lead to increased phosphorylation which could weaken the binding of the highly basic C-terminus to the ribosome. On the other hand, loss of phosphorylation would be expected to have the opposite effect. Indeed, we found that deletion of RIM15, the kinase reported to modify T607 (Dokládal et al. 2021), led to a negative genetic interaction with *rei1Δ reh1Δ*. However, single point mutations at positions 607 and 632 designed to act as phosphomimetic mutants did not suppress *rei1Δ reh1Δ*. Consequently, it remains an open question whether or not Lsg1 is a substrate of Ppq1 and whether or not phosphorylation of its C-terminus modulates its association with 60S.

Although the yeast genome contains both *REI1* and *REH1*, which share a redundant function, these two genes have also diverged in function. Only Rei1 appears to be involved in the release of Arx1. Whereas both proteins bind to the same sites on the ribosome, Rei1 precedes Reh1 in binding. We have recently shown that Reh1 is the last assembly factor to be released from the nascent 60S (Musalgaonkar et al. 2025) and that it is released in early rounds of translation elongation, after subunit joining. Unlike other ribosome assembly factors, including Rei1, the expression of Reh1 correlates with factors involved in quality control and protein degradation and not with ribosome biogenesis (Chitale et al. 2025). Indeed, we have recently shown that Reh1 is a quality control factor that monitors defective ribosomes carrying a mutation in a critical loop of uL16 in the P site of the ribosome (Chitale et al. 2025). Thus, in yeast Rei1 and Reh1 have a common redundant function in the loading of eL24 but have diverged in functions with Rei1 acting as a bona fide assembly factor and Reh1 acquiring a role in ribosome quality control. Considering that most eukaryotic organisms contain a single gene, *ZNF622* in humans, an outstanding question is whether or not *ZNF622* combines the quality control function of Reh1 with the assembly function of Rei1 or whether separate machinery remains to be identified in human cells for quality control. The recent identification of *ZNF574* from humans, which is necessary for quality control of ribosomes with defective uL16 may hint at a more complex scenario for quality control pathways in human cells (Akers et al. 2024).

## Statements

### Data Availability

All data are contained within the manuscript.

## Acknowledgements

This work was supported by NIH grant GM127127 to A.W.J. The authors thank J. Risdal for help with cloning.

## Funding and additional information

This work was funded by NIH grant R35 GM127127 to A.W.J. "The content is solely the responsibility of the authors and does not necessarily represent the official views of the National Institutes of Health."

## Supplementary figure legend

**Figure S1.**
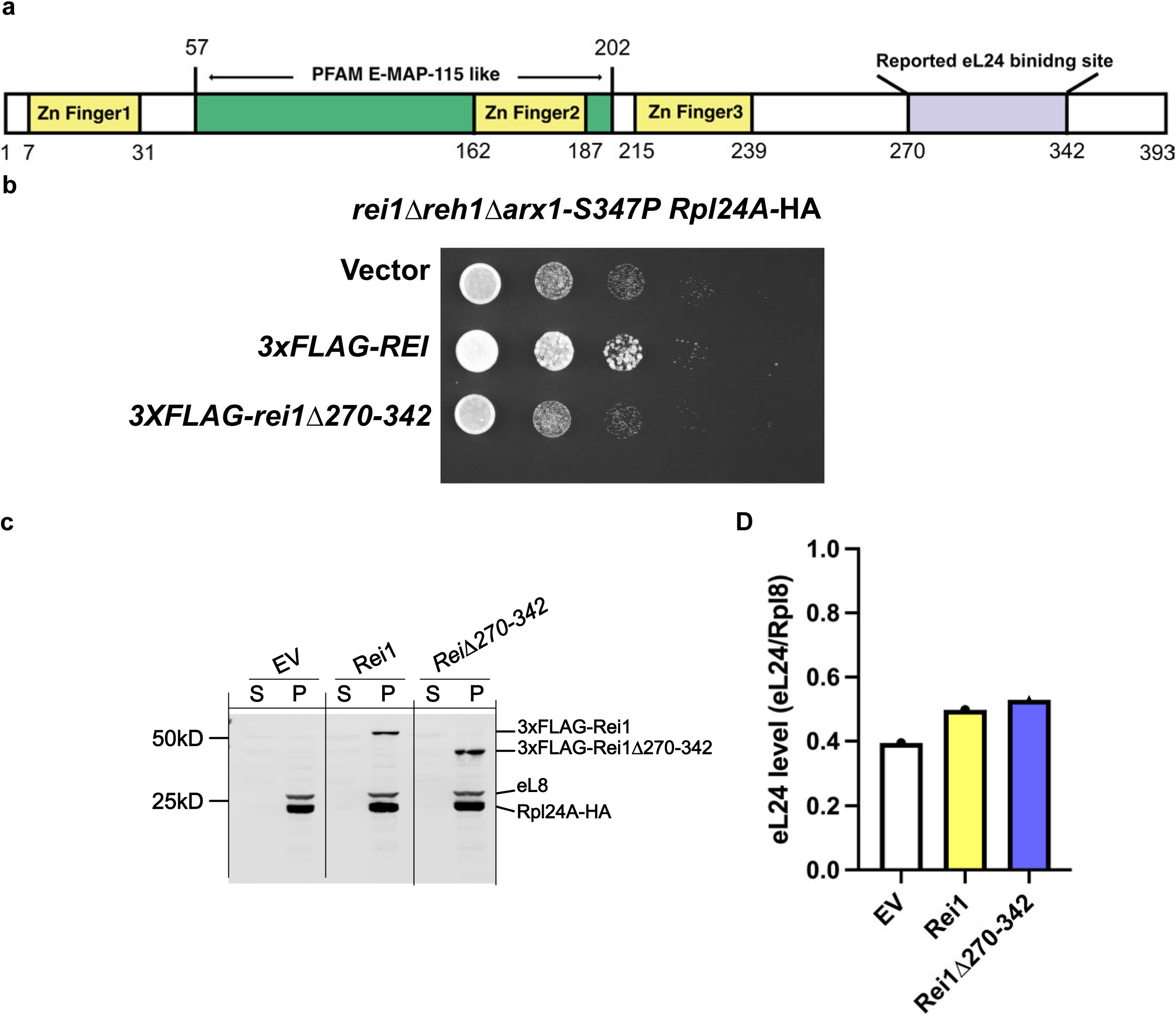
A) A schematic of Rei1 domain organization. B) rei1Δ270-342 is not functional. Serial dilutions of *rei1Δ reh1Δ arx1-S347P* strains transformed with an empty vector, a 3XFLAG tagged Rei1 plasmid, or a 3XFLAG tagged Rei1Δ270-342 plasmid. C) Western blot analysis of yeast extracts from *rei1Δ reh1Δ arx1-S347P* strains transformed with an empty vector, a 3XFLAG tagged Rei1 plasmid, or a 3XFLAG tagged Rei1Δ270-342 plasmid, following a sucrose cushion assay. Supernatant(S) or pellet(P) were collected and probed with anti-FLAG (for Rei1) and anti-HA (for Rpl24A) and Rpl8 antibodies. D) Quantification of eL24 level in rei1Δ reh1Δ arx1-S347P strains. Rei1 signal was normalized to Rpl8 to account for ribosome loading. Comparable levels of eL24 associated with the ribosome in the presence of Rei1 or Rei1Δ270-342 suggest that Rei1 270-342 is not the binding site for eL24.

**Figure S2.**
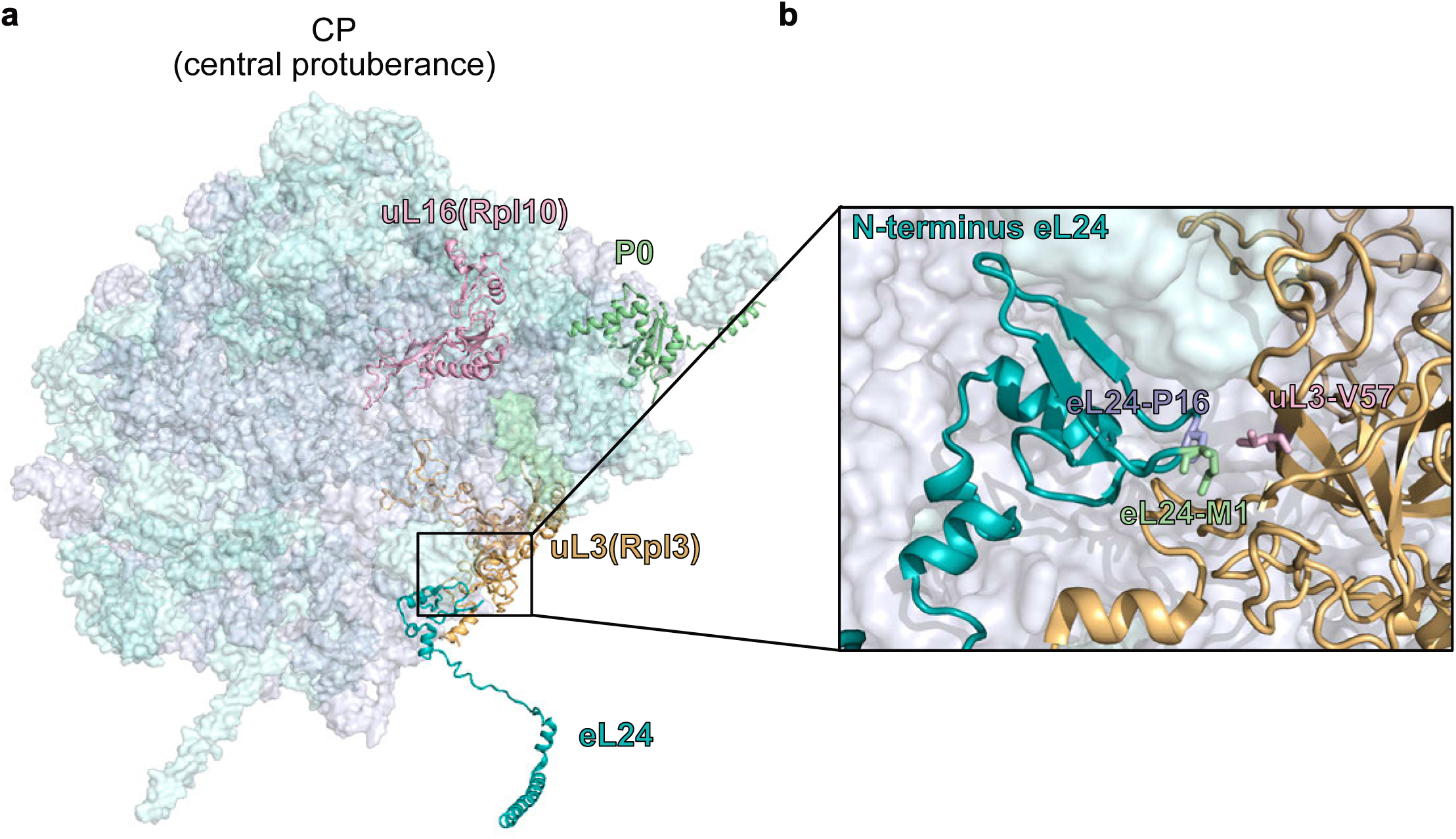
A) Surface representation of the yeast 60S ribosomal subunit (PDB: 4V88). Key ribosomal proteins are highlighted in cartoon representation: eL24 (teal), uL3/Rpl3 (light orange), uL16/Rpl10 (pink), and P0 (light green). B) Close-up view of the 60S subunit highlighting the N-terminus of eL24 and the suppressor residue V57 of uL3.

**Figure S3.**
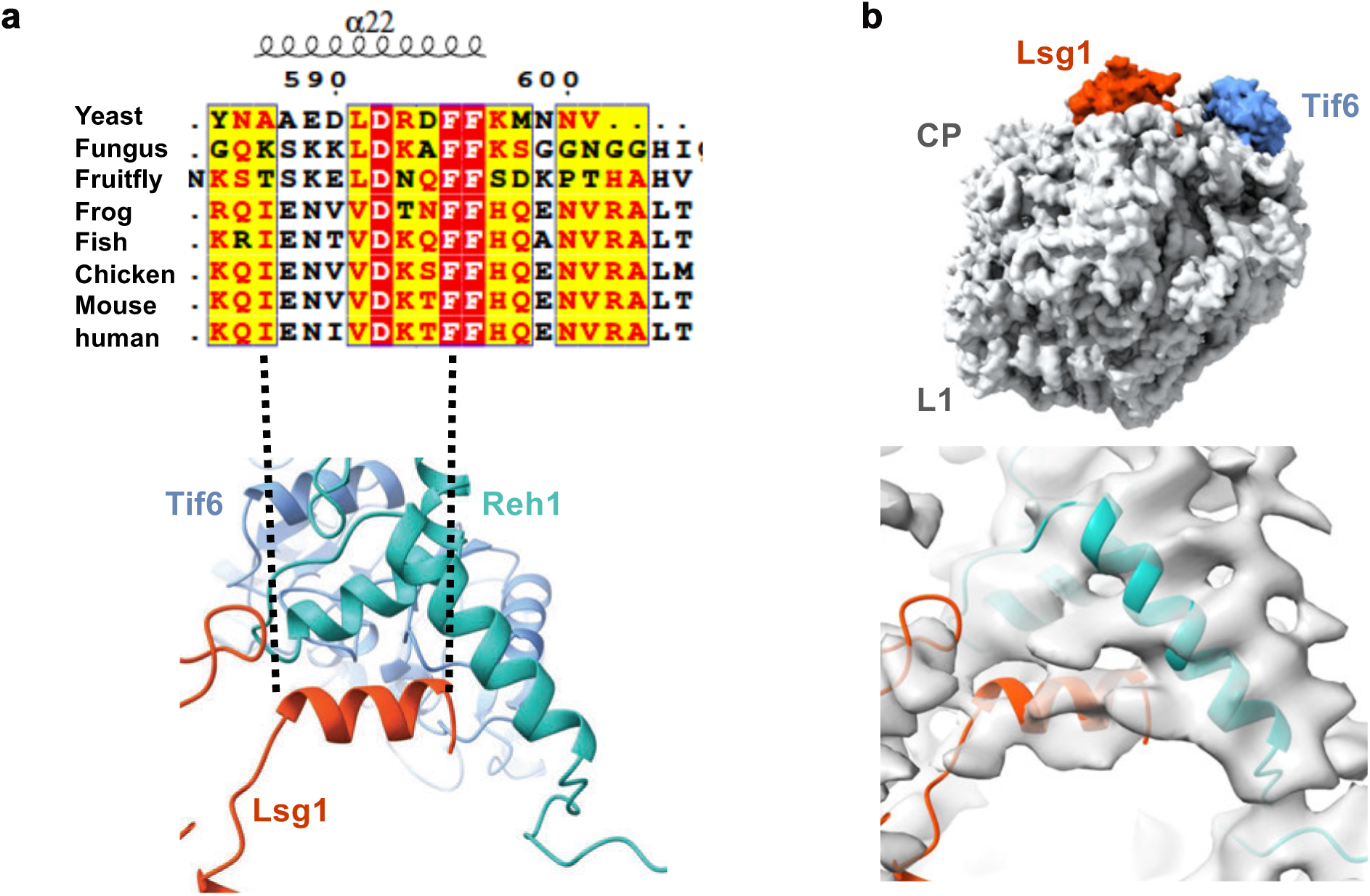
A) Multi-sequence alignment of Lsg1 and its homologs. Sequences were aligned using T-coffee, then output using ESPript 3.0. Aligned sequences: protein, organism (access No.): Lsg1, *S. cerevisiae S288C* (NP_011416.3); Large subunit GTPase 1 homolog, *N.crassa* (KAK3952141.1); Large subunit GTPase 1 homolog, *D. melanogaster* (Q9W590.1); large subunit GTPase 1 homolog, *X. tropicalis* (XP_002937880.1); Lsg1 *D.rerio*, (NP_997807.1); large subunit GTPase 1 homolog, *G. gallus*, (Q5ZJD3.1); large subunit GTPase 1 homolog isoform 1, *M. musculus* (NP_835170.1); large subunit GTPase 1 homolog, *H. sapiens* (NP_060855.2). The alignment highlights the conserved α22 helix region and is mapped onto the corresponding AlphaFold structural cartoon model with Tif6 (light blue), Reh1 (cyan) and Lsg1 (red). B) Cryo-EM density of the cytoplasmic pre-60S particle obtained in this study (top) and a zoomed-in view of the region encompassing the Lsg1 interaction interface (bottom).

**Figure S4.**
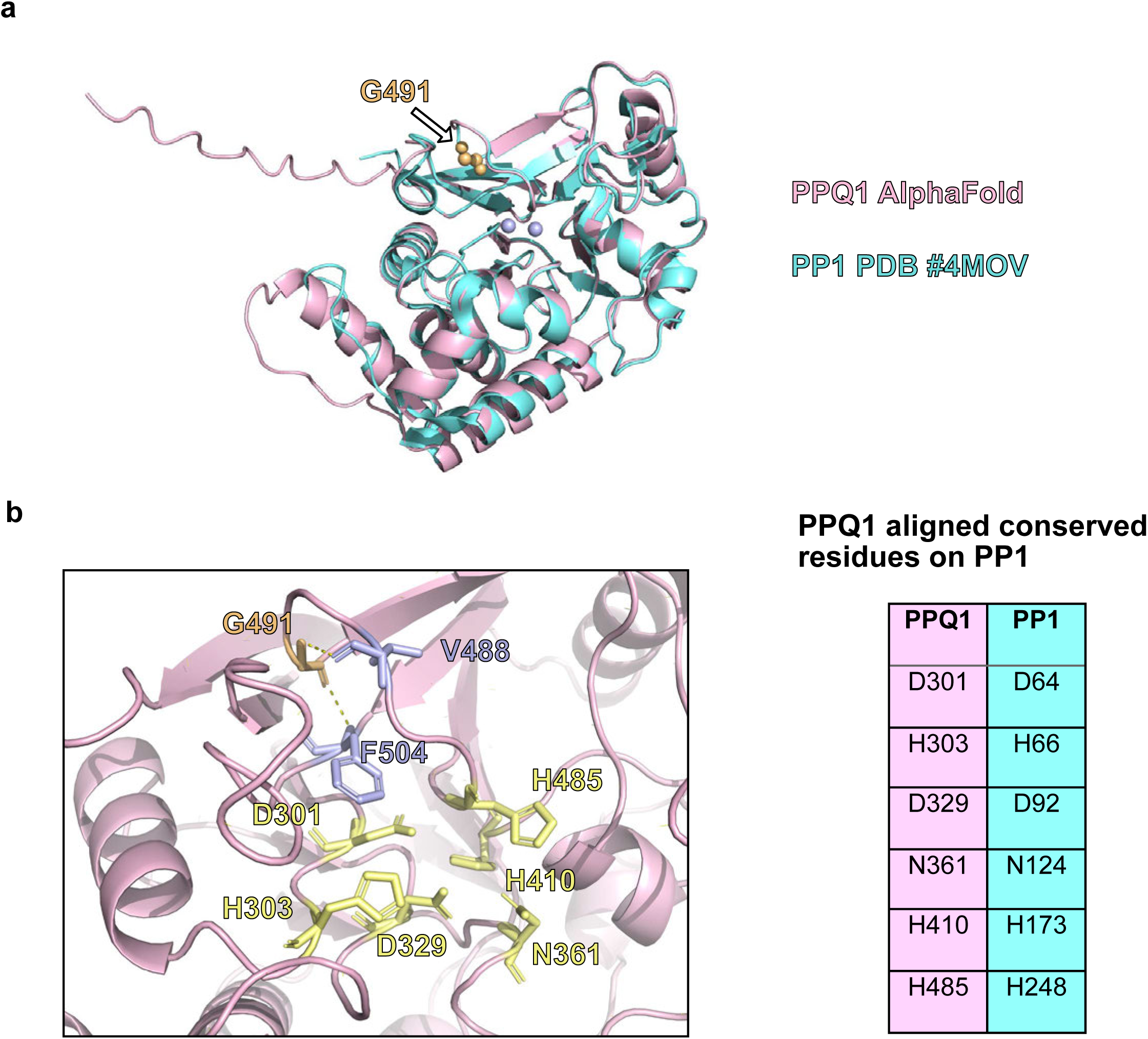
AlphaFold3 predicted Ppq1 catalytical center aligned with Pp1 well. A) Ppq1 (aa238-549) structure was predicted by AlphaFold3 (pink) and aligned with Pp1 (cyan, PDB 4MOV) in cartoon model. G491 residue was shown in sphere and colored by orange. B) Closer view of predicted catalytical center on Ppq1. Conserved residues (D301, H303, D329,N361, H410, and H485)on Ppq1 correlated to Pp1 were shown in stick (yellow). Two amino acids, V488 and F504 (purple) that G491 (orange) potentially interacts with were also shown in stick.

